# Discovery of a new evolutionarily conserved short linear F-actin binding motif

**DOI:** 10.1101/2025.04.16.649135

**Authors:** Themistoklis Paraschiakos, Biao Yuan, Kostiantyn Sopelniak, Michael Hecht-Bucher, Lisa Simon, Ksenija Zonjic, Dominic Eggers, Franziska Selle, Jing Li, Stefan Linder, Thomas C. Marlovits, Sabine Windhorst

## Abstract

Regulation of the actin cytoskeleton by actin binding proteins (ABPs) is essential for cellular homeostasis, and the mode of actin binding determines the activity of ABPs. Here, we discovered a novel “Short linear F-actin binding motif (SFM)” on the basis of the cryo-EM structure of the ITPKA-F-actin complex. We developed the computational pipeline SLiMFold, which identified 103 human SFM containing-proteins exhibiting diverse cellular functions. The SFM probably developed *ex nihilo* and remained conserved in eukaryotes, with a binding affinity to F-actin ranging from 13 to 89 µM. Furthermore, we uncovered the essential amino acids of this SFM for F-actin binding and affinity modulation. Together, the SFM seems to serve as a low affinity anchor to target proteins to F-actin, in order to connect the regulation of actin dynamics with broad cellular functions. These findings will shed new light on the role of a wide variety of proteins.

## Introduction

Actin filaments (F-actin) form a dynamic cytoskeletal framework in eukaryotic cells, driving essential processes such as cell motility, division, and morphogenesis. These processes are orchestrated by different interacting proteins, referred to as actin-binding proteins (ABPs), along with signalling and scaffolding proteins.^1^ These proteins assist in polymerisation, nucleation, elongation or capping, severing, crosslinking, or bundling of actin filaments, and are required to coordinate spatially restricted actin turnover. Their impact on actin dynamics depends on the affinity of the actin-binding domain (ABD) and, in most cases, on cellular stimulation^2^. Many ABPs are inhibited in resting cells, and a lot of them are activated by small GTPases, RhoA, Cdc42, or Rac^3^. Failures in the regulation or expression of ABPs as well as mutations can lead to severe diseases, such as neuronal and immunological disorders and cancer^4–8^.

Among the ABPs, a lot of conserved domains and motifs have been identified, upon which ABP-families with similar functions were defined. Supplementary Table 1 illustrates how the remarkable variety of motifs and domains matches the diversity of cellular actin functions. Despite this diversity, a lot of ABPs bind to a hydrophobic cleft between actin subdomain 1 and 3, such as cofilin-1, T-plastin, myosin and gelsolin^9^. Here, the binding modus and/or the affinity is different, resulting in competition of the ABPs to each other in response to cellular stimulation or under certain pathological conditions^10,11^.

Alongside globular domains with properly folded three-dimensional structure, such as that of cofilin-1, T-plastin, myosin and gelsolin, unstructured regions have been shown to play an essential role in many dynamic cellular processes^12–21^. Among these, the most common functional modules are short amino acid stretches known as short linear motifs (SLiMs) ^22–25^. The abundant G-actin binding WH2 (Wiskott-Aldrich homology 2) motif is a prominent example of a SLIM, and structure elucidation revealed that it binds to G-actin at the hydrophobic cleft between actin subdomains 1 and 3, thereby modulating polymerisation and filament dynamics^26^. Missense mutations located inside the WH2 motif are associated with diseases, i.e. the human Leiomodin-3 (LMOD3), where mutations R543L^27^ or L550F^28^ are associated with Nemaline myopathy 10 (NEM10). Moreover, WH2 motifs can also be part of bacterial effector proteins, often leading to restructuring of the host cell actin cytoskeleton and promoting either efficient colonisation (VopL/VopF) or bacterial motility during infection (Sca2 and RickA)^29–32^.

In this study, we identify a novel short linear F-actin binding motif (SFM) that mediates F-actin binding in diverse human proteins with low affinity. Comparative sequence analyses indicate that the SFM is under purifying selection and acts as a low-affinity anchor, targeting various cytosolic proteins to the actin cytoskeleton. Notably, our findings not only unify previously observed yet unclassified sequences under this defined motif, but also enable the prediction of new SFM-containing proteins. These observations point to a broader role for the SFM in cytoskeletal organization under both physiological and pathophysiological conditions and may uncover new cellular actin networks and mechanisms.

## Results

### Structural insight into ITPKA in F-actin binding

A high number of actin-binding proteins (ABPs) contain uncharacterized domains responsible for F-actin binding, including the human Inositol-trisphosphate 3-kinase A (ITPKA)^10,33–36^. This InsP_3_Kinase isoform constitutively binds to F-actin, and due to homodimer formation it exhibits F-actin bundling activity^36^. Under physiological conditions, ITPKA contributes to control the postsynaptic actin architecture of hippocampal neurons. However, ITPKA is also expressed in different malignant tumor cells, and its actin bundling activity induces the formation of cellular protrusions essential for invasion of cancer cells^34,37^.

To investigate the molecular basis of ITPKA for F-actin binding in detail, we solved the cryo-EM structure of the ITPKA-F-Actin complex at 2.97 Å resolution. In this structure, the F-actin binding domain (ABD) of ITPKA is clearly visible, whereas the catalytic domain and the linker connecting it to the ABD were not resolved (Figure 1a, and Extended Data Figure 1), confirming that the C-terminal InsP_3_Kinase domain is not directly involved in F-actin binding. The amino acids Arg28 to Ala49 bind to the hydrophobic cleft between actin subdomains 1 and 3, and also contacts the DNase-binding loop of the adjacent actin subunit (A_−2_) (Figure 1a, and Extended Data Figure 1). In parallel to us, the Belyy group resolved the cryo-EM structure of F-actin bound by F-tractin, which represents the core actin binding domain of ITPKA and is used as F-actin marker for live cell imaging^11,38^. This structure matches our findings of the F-actin binding mode^11^.

**Figure 1:**
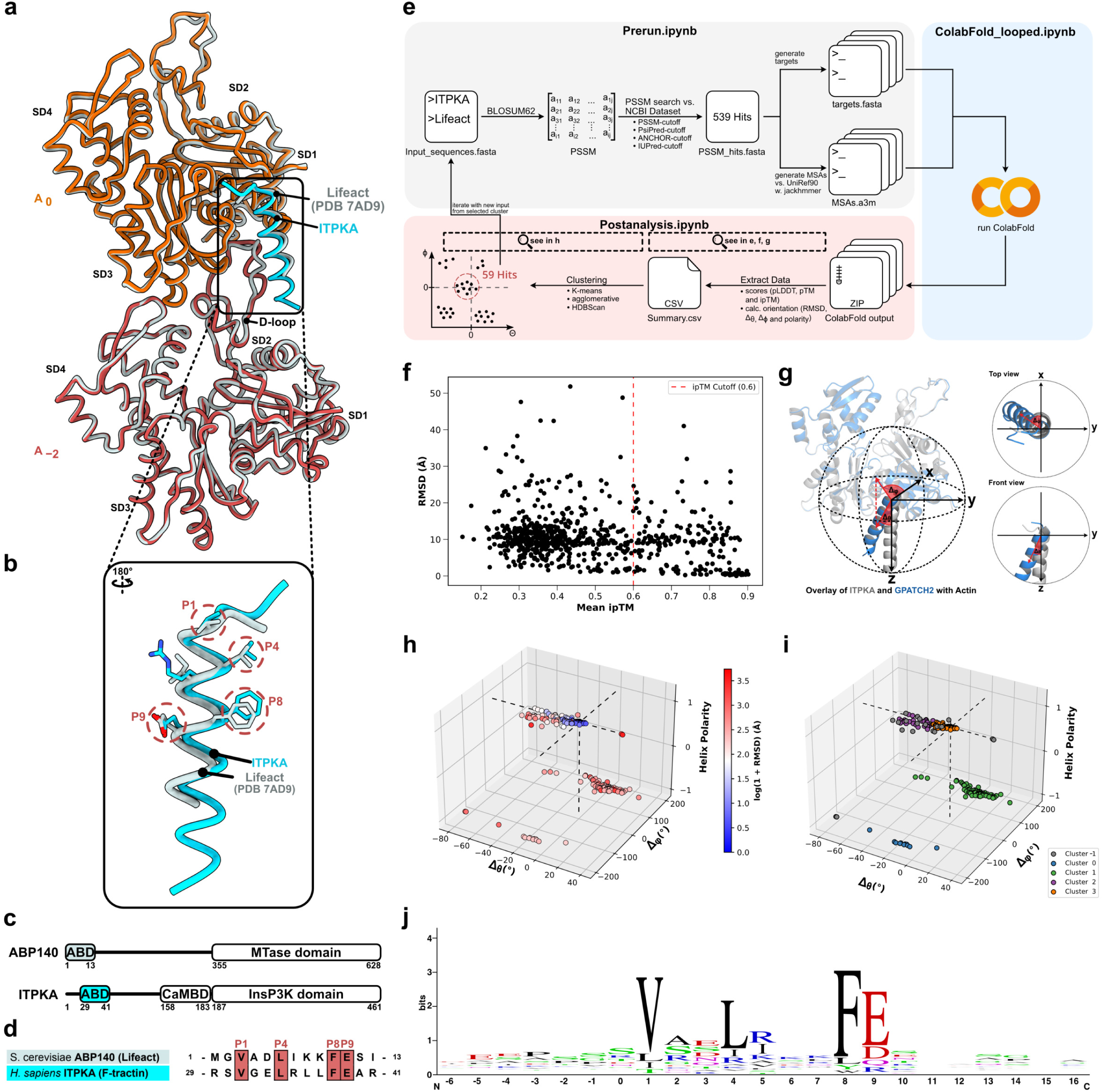
Structural alignment of ITPKA^Arg28-Ala49^ and Lifeact points towards a novel F-actin binding motif, prompting proteome-wide discovery using the SLiMFold pipeline. **(a)** Structural superimposition of the ITPKA^Arg28-Ala49^–actin complex (orange and red for the two actin subunits, cyan for ITPKA) with the Lifeact–actin structure (grey; PDB: 7AD9) shows near-identical actin conformations apart from a minor shift in the D-loop of the ITPKA-bound model. **(b)** Rotating the helices by 180° highlights the residues interacting with F-actin. Conserved side chains are represented as sticks and labeled P1, P4, P8, and P9. **(c)** Domain architecture of **S. cerevisiae** ABP140 and **H. sapiens** ITPKA. ABP140 comprises an N-terminal F-actin binding domain (ABD) and a C-terminal methyltransferase (MTase) domain. ITPKA features an N-terminal ABD, a central calmodulin-binding domain (CaMBD), and a C-terminal inositol 1,4,5-trisphosphate 3-kinase (IP3K) domain. Notably, the F-actin binding motifs, derived in this study, are located within intrinsically disordered regions, which is typical for short linear motifs (SLiMs). **(d)** Sequence comparison underscores structural and sequence conservation, conserved positions P1, P4, P8, and P9 are labeled in red. **(e)** SLiMFold pipeline overview. The SLiMFold pipeline started with the hypothesized SLiMs of Lifeact and ITPKA^Arg28-Ala49^ Sequence, on which basis a position-specific scoring matrix (PSSM) was calculated to identify motif hits from NCBI databases which were filtered by PSSM scores, IUPRED, ANCHOR, and PSIPRED. This yielded 539 hits in the first iteration. Thereafter, multiple sequence alignments (MSAs) were calculated locally using UniRef90 with modified jackhmmer filters, followed by multimer predictions on ColabFold. The predicted structures were analyzed by extracting AlphaFold2 scores, computing RMSD and angle metrics (θ, φ). These data were clustered to identify 59 new sequences that carry the suspected motif. **(f-i)** Detailed illustration of calculated metadata from (e). **(f)** Scatterplot of predicted peptide–actin complexes plotting mean ipTM vs. RMSD relative to ITPKA^Arg28-Ala49^. A red dotted line at ipTM=0.6 marks the threshold below which interactions are unreliable. **(g)** Helix orientation was assessed by comparing Δφ and Δθ angles in a polar coordinate system after superimposing each complex onto the ITPKA^Arg28-Ala49^–actin reference. **(h)** A 3D plot of Δφ, Δθ, and helix polarity (1 = same direction, −1 = opposite) maps each predicted peptide, with data points colored by log(1+RMSD). Angles near zero indicate an orientation similar to that of ITPKA^Arg28-Ala49^. **(i)** HDBSCAN clustering of these data points groups peptides based on orientation and RMSD. Outliers are assigned to Cluster –1, while Cluster 3 contains helices most closely matching ITPKA^Arg28-Ala49^. **(j)** Sequence logo depicting positional conservation among the 59 identified peptides confirms robust conservation at positions P1, P4, P8, and P9, strongest at P1 and P8.

Collectively, we found that the core actin binding domain of ITPKA only comprises 21 amino acids folded into an alpha helix. Interestingly, this peptide exhibits the characteristics of a short linear motif (SLiM), and its structure resembles that of the F-actin probe Lifeact^11,39,40^. Based on this consideration, we performed a structural alignment of ITPKA^Arg28-Ala49^ with Lifeact (PDB: 7BTE and 7AD9) and found highly similar helix positioning, including the aligned positions Val (P1), Leu (P4), Phe (P8), and Glu (P9) (Figure 1b,c, d).

This discovery let us to propose the F-actin binding SLiM VxxLxxxFE, and to determine whether other human proteins include this motif, we developed a computational pipeline, called SLiMFold (Figure 1e). First, we derived a Position Specific Scoring Matrix (PSSM) from the aligned sequences of ITPKA^Arg28-Ala49^–and Lifeact (see Figure 1e and Extended Data Figure 2), and used it to screen against the NCBI human protein database. Next to this PSSM cutoff, we applied PSIPRED^41^, ANCHOR^42^, and IUPred^43^ cutoffs to highlight candidates containing this SLiM (Figure 1e upper panel). For each of these 539 hits a multiple sequence alignment (MSA) was performed via jackhmmer^44^ against UniRef90 database, with optimized parameters to improve both speed and coverage.

Once the MSAs were compiled, we ran a custom ColabFold notebook to predict the structure of various peptide–actin complexes, leveraging the user-provided MSAs instead of recalculating new ones. Each peptide–actin model was scored by pLDDT, pTM, and ipTM, and any candidate with mean ipTM below 0.6 (averaged over the top three ranked models) was excluded (Figure 1e lower panel, and Figure 1f).

To assess structural convergence with the reference ITPKA^Arg28-Ala49^ motif, each candidate underwent a detailed comparison against the ITPKA^Arg28-Ala49^–actin complex. We measured RMSD for α-carbon atoms (positions P1–P9), helix orientation (Δθ, Δφ) using a polar coordinate system, and helix polarity (Figure 1f, g, h). The resulting models were then clustered using HDBSCAN^45^, which effectively segregates core clusters from outliers (Figure 1i). Cluster 3—enriched in 59 gene sequences—displayed minimal RMSD and orientation shifts relative to ITPKA, suggesting a highly similar helix arrangement for actin binding (Figure 1i). A sequence logo of these 59 clustered hits confirmed strong positional conservation at P1, P4, P8, and P9, with P1 and P8 showing the highest degree of conservation (Figure 1j). Since this actin binding motif belongs to the SLiMs, it was designated as Short linear F-actin binding Motif (SFM).

### Validation of SFM-peptides *in cellulo* and *in vitro*

To validate binding of the predicted SFM-peptides to F-actin, we randomly selected candidates, and split them into two groups. Group 1 included the RhoGTPase-guanine exchange factors (GEFs) FGD1, FGD4, DENNDC1, and ARHGEF11, as well as DIXDC1, a protein that is involved in regulation of wnt-signalling (Figure 2a). Group 2 comprised the actin bundling proteins SHROOM 3, ESPNL, and PPP1R9A, the ubiquitin specific protease USP54, the z-disc protein CEFIP and the tight junction protein CGNL1 (Figure 3a). Group 1 candidates were validated by F-actin-colocalization with EGFP-peptides or full-length proteins in primary human macrophages as well as in H1299 lung cancer cells, and Group 2 candidates were validated by F-actin co-sedimentation assays with recombinant peptides.

**Figure 2:**
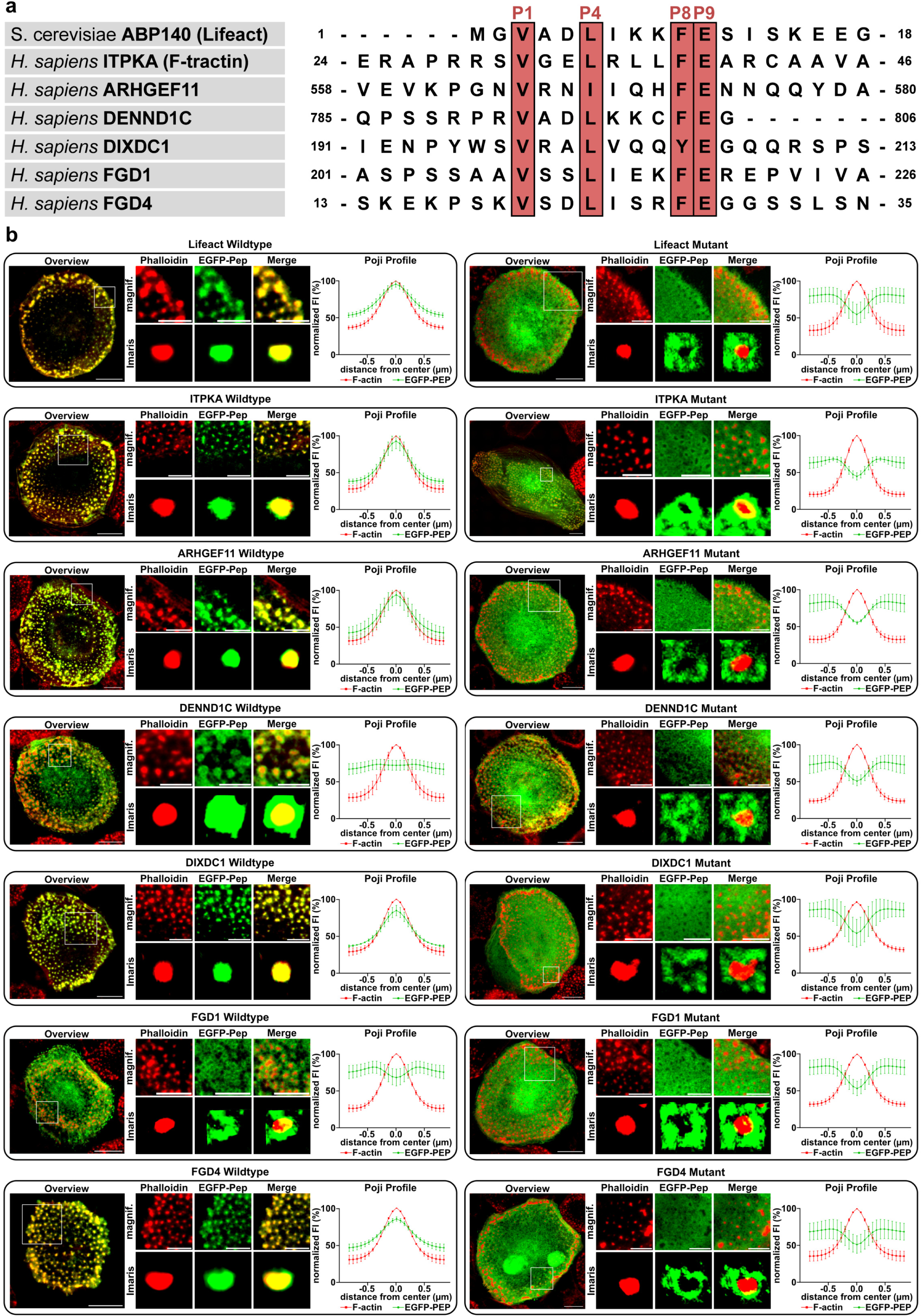
Validation and functional analysis of the novel short linear F-actin binding motif (SFM) by cellular F-actin colocalization. **(a)** Five peptides identified by the SLiMFold pipeline were aligned with the sequences of ITPKA and Lifeact; the conserved positions P1, P4, P8, P9 are highlighted in red. To probe their functional significance, the peptides were cloned into an pEGFP vector, and alanine mutations were introduced at P1 and P8. **(b)** Primary human macrophages were transfected with the vectors coding for the EGFP-tagged peptides and stained with phalloidin-568 to visualize F-actin (red). Representative cell images are shown including zoomed-into one podosome substructure, scale bars: 10 and 5 µm, respectively. Colocalization is evident when EGFP-tagged peptides (green) overlap with phalloidin-stained actin (red). Right panels: Imaris 3D reconstructions and Poji radial profiles confirm varying degrees of peptide–actin colocalization. Shown are Poji radial profiles of fluorescence intensities, at a z plane of highest F-actin intensity, with the mean values of F-actin and respective peptide ± standard deviation of at least 3 cells, and 240 podosomes per transfected peptide. Poji radial profiles were normalized to set the intensity values from 0% to 100%.

**Figure 3:**
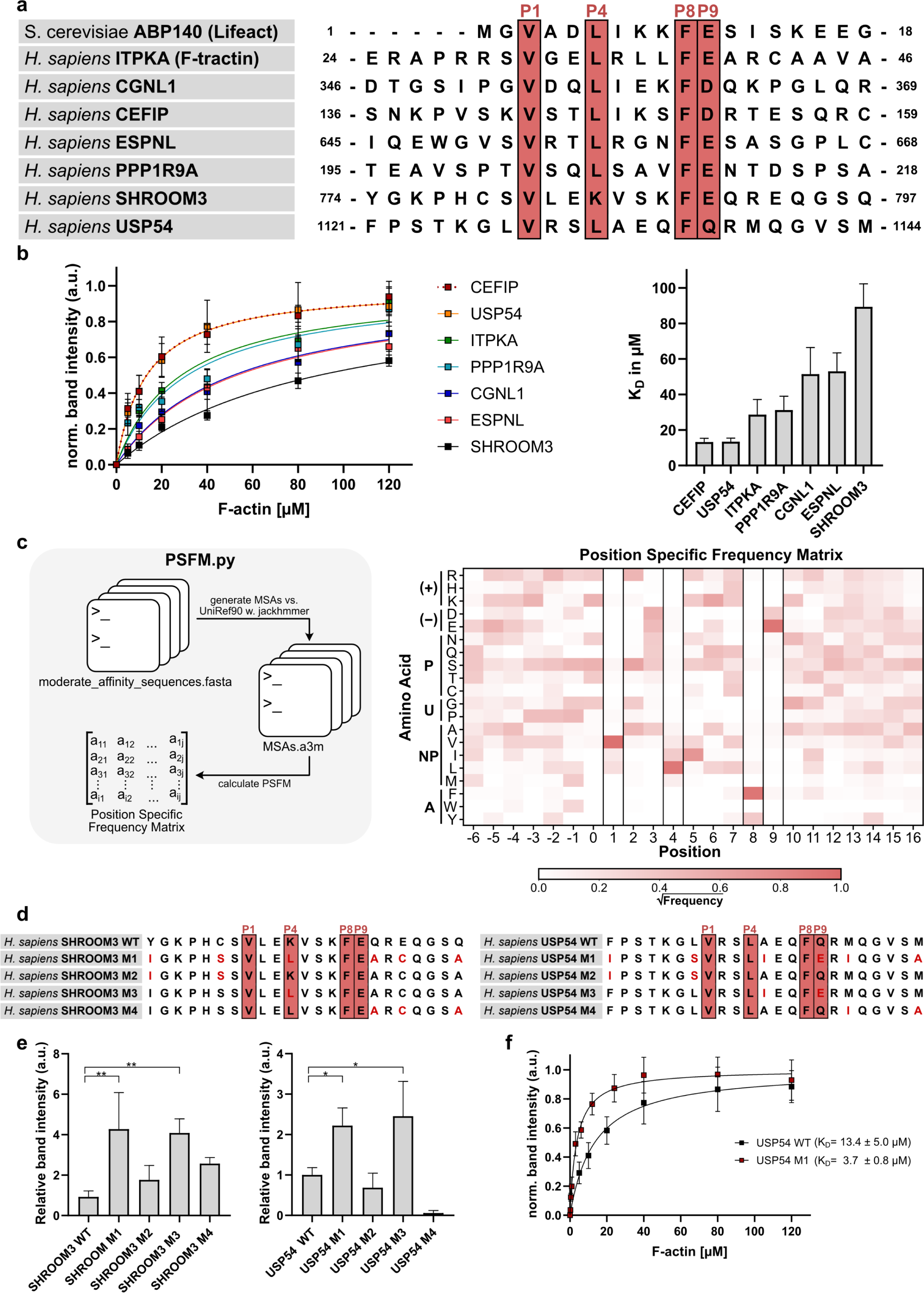
Determination of SFM-binding affinities to F-actin, and identification of affinity-modulating amino acids. **(a)** Six further peptides identified by the SLiMFold pipeline were aligned with the sequences of ITPKA and Lifeact, and the conserved positions P1, P4, P8, and P9 highlighting in red. The peptides were expressed in bacteria, and after purification F-actin pull-downs were performed. Binding of the peptides were analyzed by Western blotting, and to calculate relative dissociation constants (K_D_ values), the F-actin concentrations were plotted against normalized band intensities. The right panel provides the K_D_-values for all peptides tested. **(c)** In order to identify the amino acids modulating the affinity of the peptides to F-actin, their frequency was analyzed by generating a position specific matrix (PSFM) using MSA from CEFIP, USP54, ITPKA, PPP1R9A, Lifeact, ARHGEF11, FGD4, DIXDC1. Y-axis represents positively charged (+), negatively charged (-), polar (P), unique (U), non-polar (NP) and aromatic (A) residues, and X-axis the position inside the SFM. This matrix, color-coded by √(amino acid frequency), pinpointed less (white) and more frequent (red) residues. **(d)** SHROOM3 and USP54 wild-type (WT) sequences and corresponding mutants, including the key residues P1, P4, P8, P9. M2 features mutations N-terminal to the motif, M3 contains mutations within the motif, and M4 includes mutations C-terminal to the motif. M1 incorporates all mutations (N-terminal, within, and C-terminal). Newly mutated positions are labeled in red. **(e)** Quantified relative band intensities from SHROOM3 and USP54 pulldown assays, normalized and averaged across replicates, highlighting differences in binding affinities between WT and mutant peptides. (f) For USP54 M1 addiotinally the affinity to F-actin was determined. Statistical significance is denoted by asterisks: *\*p < 0.05*; *\*\*p < 0.01*.

The Group 1 SFM peptides, including the positive controls ITPKA^Arg28-Ala49^ and Lifeact, were transiently expressed as EGFP-fusion peptides in primary human macrophages, allowing a precise quantification of co-localization with F-actin subsets^46^. We additionally tested site-specific mutants (V→A and F→A) at positions 1 and 8. As shown in Figure 2 (and Extended Data Figure 4), Lifeact, ITPKA^Arg28-Ala49^, FGD4 and ARHGEF11 showed prominent co-localization with podosome cores. However, DENND1C mainly localized to the podosome cap, and also FGD1 showed a broader distribution, encompassing parts of the podosome core. On the other hand, all P1/P8 mutants were diffusely distributed, confirming that positions P1 and P8 inside the SFM are essential for F-actin binding. Similar results were obtained also in H1299 lung cancer cells (Extended Data Figure 5). Together, these data indicate that the SFM binds to F-actin with different efficiency, and preference to F-actin subpopulations.

To further confirm these findings, we tested full-length versions of selected proteins in H1299 cells and introduced the mutations at P1/P8 position. Colocalization with F-actin was observed for the wild-type full-length proteins but completely disappeared upon mutation of the residues at P1/P8 position (Extended Data Figure 6). This result highlights the critical role of the SFM in mediating F-actin binding, not only in isolated peptides but also in full-length proteins.

For Group 2 candidates affinities of peptides to F-actin were tested. For this purpose, the peptides were recombinantly expressed as EGFP-fusion peptides in bacteria, and after purification its binding to F-actin was assessed by a co-sedimentation assay. After employing different F-actin concentration, dissociation constants (K_D_) were calculated, showing that the SFMs of CEFIP and USP54, of ITPKA and PPP1R9A, and of CGNL1 and ESPNL bound with very similar affinities to F-actin (13 µM, 30 µM, and 50 µM) while the affinity of SHROOM 3 was lowest (90 µM) (Figure 3). These data reveal a relative low F-actin binding affinity of the SFM-peptide with high variations among the different peptides.

### Amino acids in between the conserved Valine (P1) and Phenylalanine (P8) determine affinity of the SFMs

Our data depicted in Figure 3b reveal that although all SFM-peptides tested included the conserved amino acids at P1/P8 position, their binding affinities to F-actin were different. This raises the question as to whether amino acids (aa) flanking the motif modulate the affinity.

To address this, we aimed to identify the aa inside the SFM mediating the highest affinity to F-actin. For this purpose, the jackhammer generated MSAs of the peptide sequences from ITPKA, CEFIP, PPP1R9A, USP54, FGD4, ARHGEF11, DIXDC1, and Lifeact were used to generate a position-specific frequency matrix (PSFM) (Figure 3c).

Aligning homologous sequences from these motifs served as a “contrast enhancer”, revealing positions that appear intolerant to certain physiochemically similar amino acid groups (white squares in Figure 4c) or potentially essential for F-actin binding (red squares in Figure 4c). On the basis of this matrix, three hotspots were defined, an N-terminal, an internal, and a C-terminal motif. To test whether substitution of low frequency amino acid with high frequency amino acid would enhance the binding affinity to F-actin, targeted mutations in SHROOM3 and USP54 were introduced according to these hotspots. In SHROOM3, point substitutions Y1I/C6S (M2), K11L (M3), and Q17A/E19C/Q23A (M4) were produced, along with a triple-hotspot variant (M1). Similarly, USP54 mutants F1I/L7S (M2), A12I/Q16E (M3), M18I/M23A

**Figure 4:**
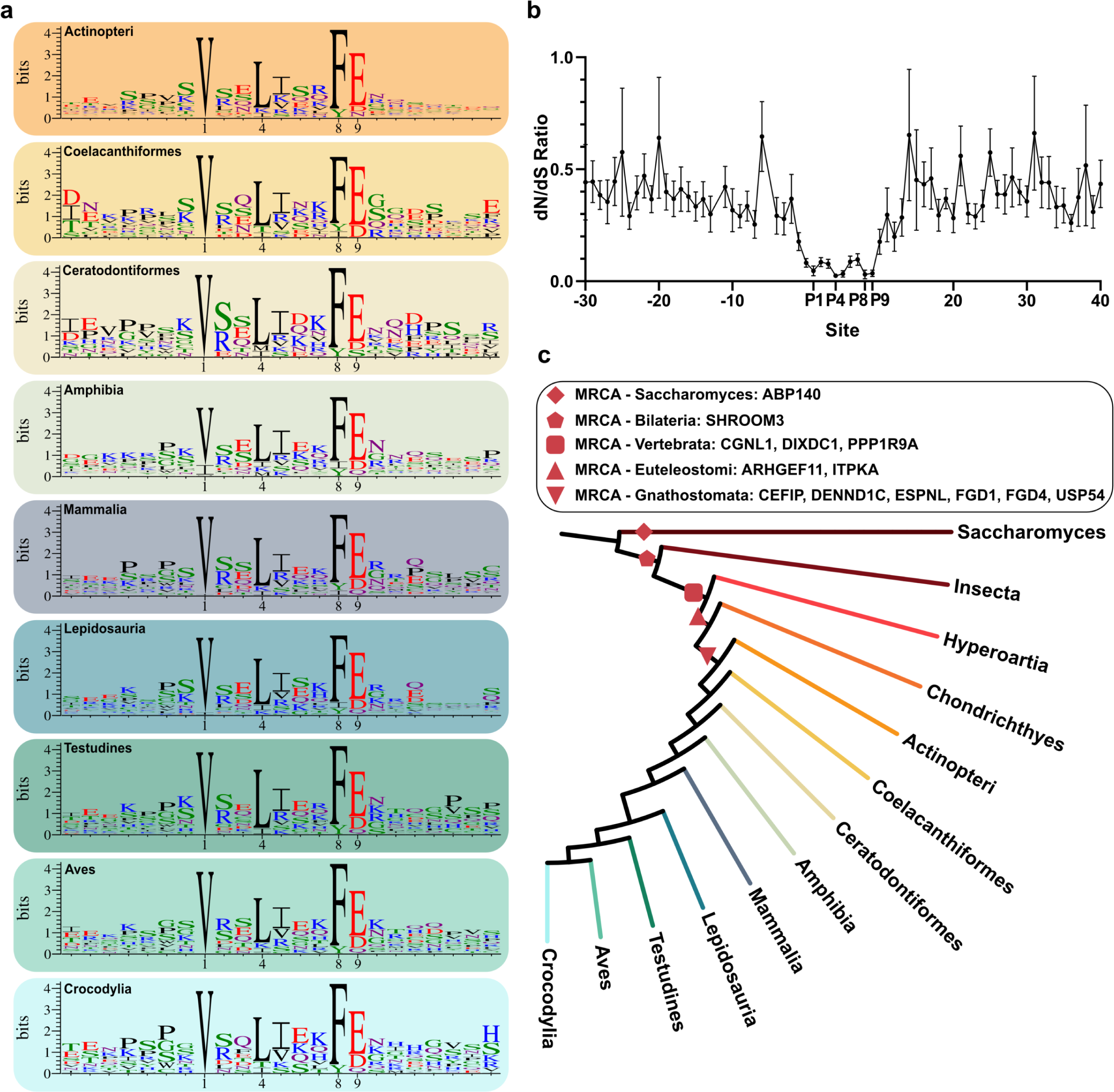
Evolutionary analysis of validated SFM-containing proteins. **(a)** Sequence logos of SFMs from representative organism classes (e.g., *Actinopteri, Coelacanthiformes*, *Amphibia*, *Mammalia*, *Lepidosauria*, *Testudines*, *Aves*, and *Crocodylia*). Key motif positions P1, P4, P8, and P9 are labeled. Larger letters indicate higher residue conservation, reinforcing the importance of these positions for F-actin binding. **(b)** A dN/dS ratio analysis of all validated SFM-containing peptides is plotted ± SEM, with amino acid position on the x-axis and the dN/dS ratio on the y-axis. The conserved SFM residues (P1, P4, P8, P9) are annotated to highlight their selective pressures. Notably, these positions exhibit lower dN/dS values, suggesting strong purifying selection. **(c)** Phylogenetic tree illustrating the distribution of validated SFM-containing proteins across various species. Symbols mark the most recent common ancestor (MRCA) *Saccharomyces* (ABP140), *Bilateria* (SHROOM3), *Vertebrata* (CGNL1, DIXDC1, FGD4, PPP1R9A), *Euteleostomi* (ARHGEF11, ITPKA), and *Gnathostomata* (CEFI, DENND1C, ESPLN, FGD1, FGD4, USP54).

Indeed, after assessing binding of the wt and mutant peptides to F-actin we found that mutation of the amino acids between the conserved V(P1)/F(P8) positions (internal motif) increased binding to F-actin four-fold in SHROOM 3 (mutant S2) and three-fold in USP54 (Mutant U2) compared to wt protein. However, mutations N-or C-terminal to the V(P1)/F(P8) positions (Mutants S1, S3, U1, U3) had no significant effects (Figure 3e). Moreover, determination of USP54 M1 (triple-hotspot variant) affinity to F-actin revealed a K_D_ of 3.7 µM (Figure 3f), confirming increased binding of the mutant to F-actin.

In summary, our data show that the conserved amino acids at P1 and P8 position are required for binding of the SFM-motif to F-actin, and reveal that those residues in between modulate affinity.

### The amino acids inside the SFM are conserved over evolution

To examine the evolutionary conservation of the SFM motif, we generated sequence logos for 12 validated SFM-containing proteins across *Crocodylia*, *Aves*, *Testudines*, *Lepidosauria*, *Mammalia*, *Amphibia*, *Coelacanthiformes*, and *Actinopteri*. These seven classes were chosen because, with the exception of Lifeact (which is found only in *Saccharomyces* and thus not represented in these classes), they cover all known occurrences of the SFM in our studied dataset. The resulting logos consistently highlighted Val (P1), Leu (P4), Phe (P8), and Glu (P9) as predominantly invariant across the examined taxa. Although a small number of substitutions were observed at these positions in certain species, these core residues remained largely unchanged, indicating strong evolutionary constraints (Figure 4a). This observation aligns with our initial comparison of ITPKA and Lifeact (Figure 1), where a similar set of core residues were conserved. To accurately quantify the selective forces acting on these conserved positions, we calculated the non-synonymous to synonymous substitution ratio (dN/dS) within each SFM-containing gene across the taxa in which the motif was confidently detected. We then averaged these values around the SFM’s core (Val (P1) to Glu (P9)) for all proteins. The resulting low dN/dS ratio (relative to nearby flanking regions) strongly suggests that the SFM is under purifying (negative) selection, consistent with its implied functional importance (Figure 4b). Consistent with AlphaMissense pathogenicity scores^47^ (Extended Data Figure 8), this highlights the critical necessity of these amino acids for cell survival.

To get further insight into evolutionary development of the motif, phylogenetic trees were constructed and traced each validated SFM-containing protein back to its most recent common ancestor (MRCA). This analysis revealed a relative early appearance of SHROOM3 in *Bilateria*, followed by PPP1R9A, DIXDC1 and CGNL1 in *Vertebrata*. CEFIP and ITPKA emerged in *Euteleostomi*, and by the time of *Gnathostomata* USP54, FGD1, FGD4, ESPN, DENND1C were present (Figure 4c). Despite this broad taxonomic distribution, we found no evidence of exon duplication or shuffling that might have repurposed an older motif. These observations support a convergent evolution and a spontaneous origin (*ex nihilo*) of the SFM.

To conclude, phylogenetic analysis confirms the conserved nature of the SFM motif, and indicates an *ex nihilo* development.

### Final iteration of the SLiMFold pipeline delivers 103 new SFM-candidates with different biological roles

Finally, we refined the PSSM of the SLiMFold pipeline by incorporating our newly identified SFM peptides, then performed two additional iterations to compile a definitive list of putative human SFM proteins (Figure 5a and Extended Data Figure 9). In total, 124 final SFM sequences emerged in 103 genes (summarized in Table 1). Of these, 26 proteins—including both our experimental validations and those reported in the literature^48–68^—have been tested using truncation-based approaches for the SFM. Additionally, 16 other proteins are known to bind F-actin through more indirect methods (e.g., colocalization or co-sedimentation of the full-length protein)^69–85^. Further testing and more detailed truncation studies are still needed to validate the remaining predicted proteins.

**Figure 5:**
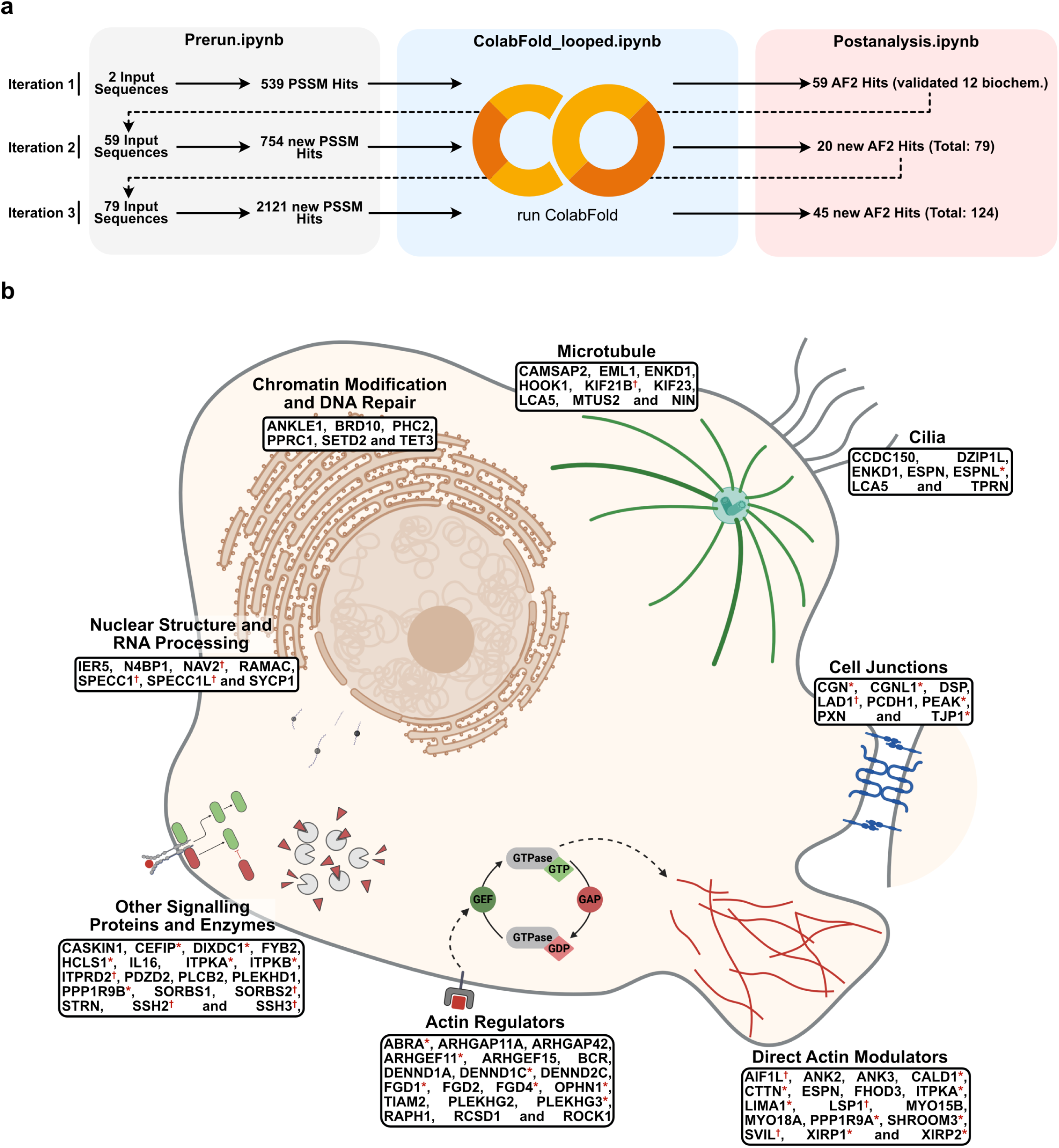
Iterative SLiMFold pipeline expansion and functional classification of newly identified SFMf-containing proteins. **(a)** Scheme of three iterative SLiMFold rounds (for detailed description, see Figure 3), including *in vitro* and *in cellulo* validation, yielding 124 peptide-sequences from 103 different genes. **(b)** Cartoon representation of a eukaryotic cell highlighting the functional categories of new SFM-bearing proteins. SFM candidates are grouped according to their principal roles and subcellular localizations, encompassing direct actin modulators, actin regulators, other signaling proteins and enzymes, nuclear structure and RNA processing, chromatin modification and DNA repair, microtubule-associated factors, cilia, and cell-junction-associated proteins. Proteins for which F-actin binding has been attributed to the SFM (e.g., validated by truncation studies) are marked with an asterisk (*), whereas proteins that bind F-actin but lack direct evidence linking the interaction specifically to the SFM are marked with a dagger (†). The cartoon was created with BioRender.com.

**Table 1.**
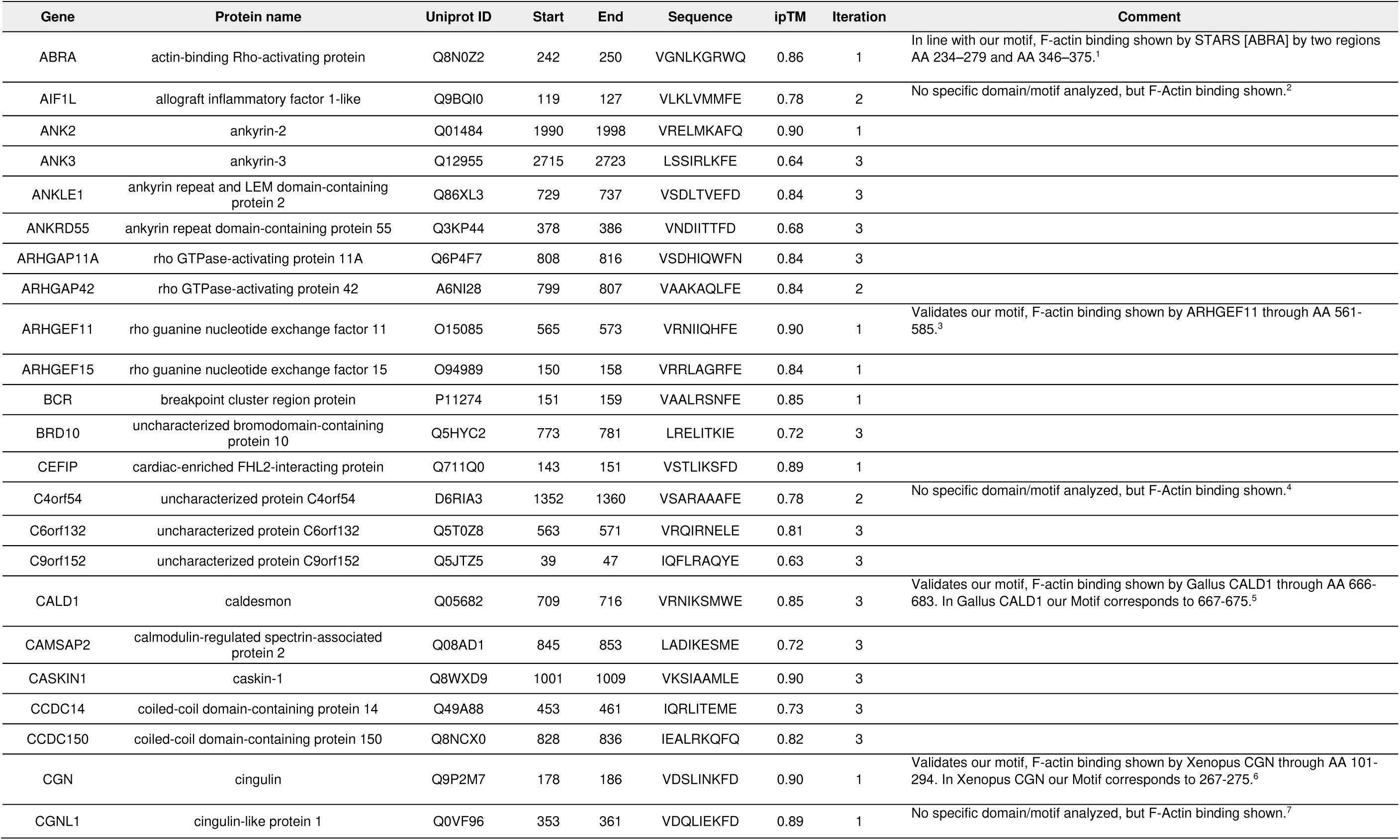
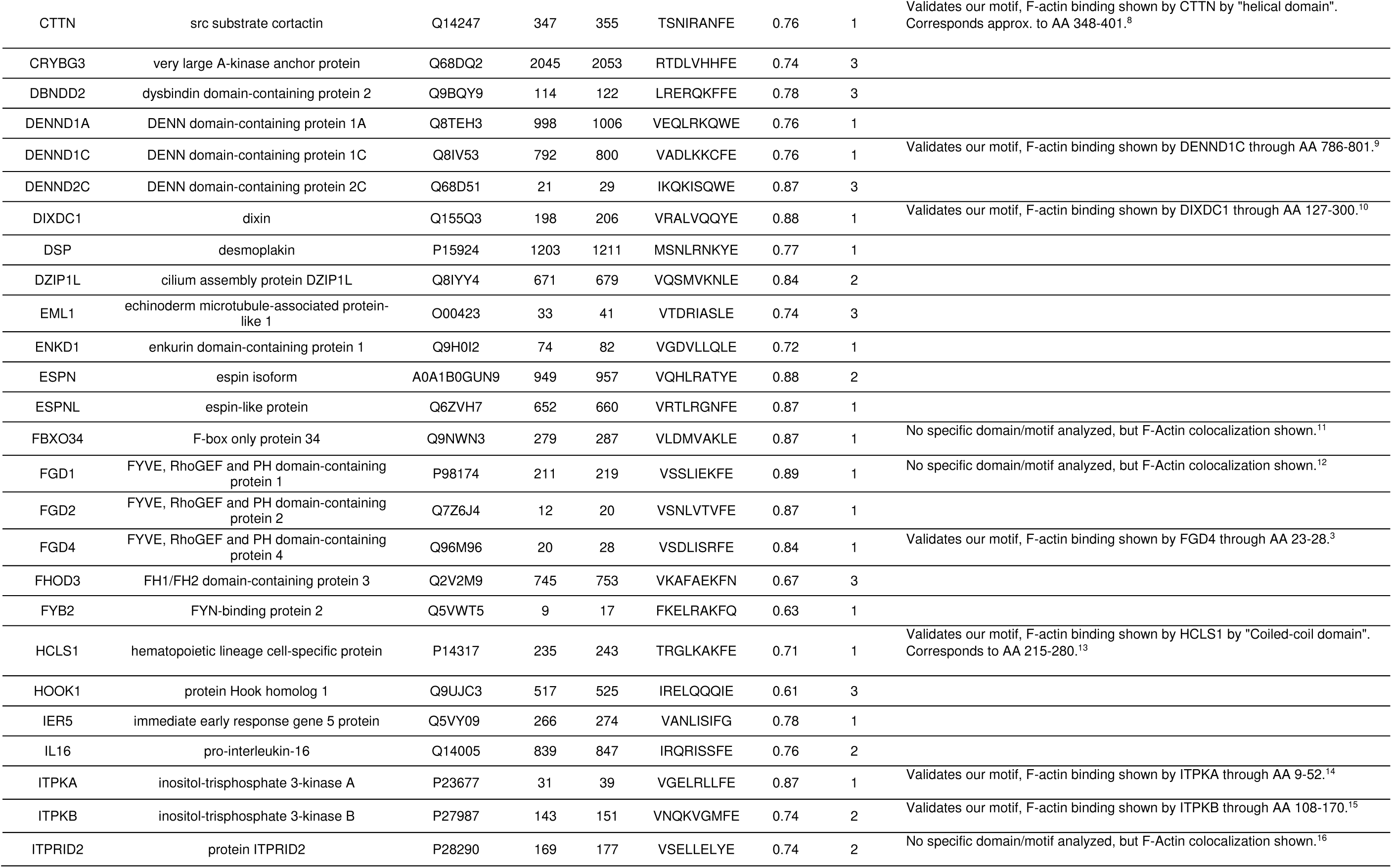
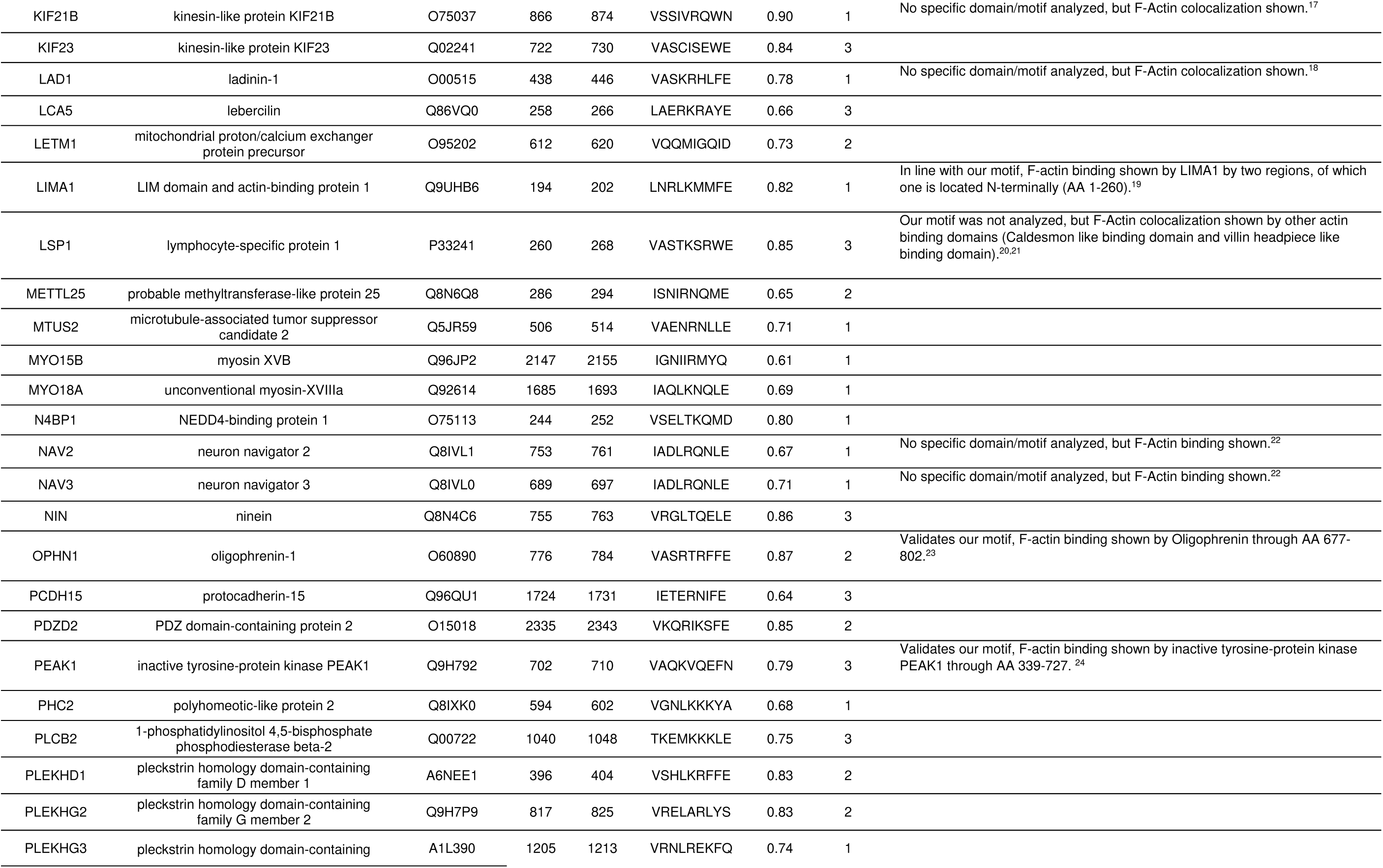
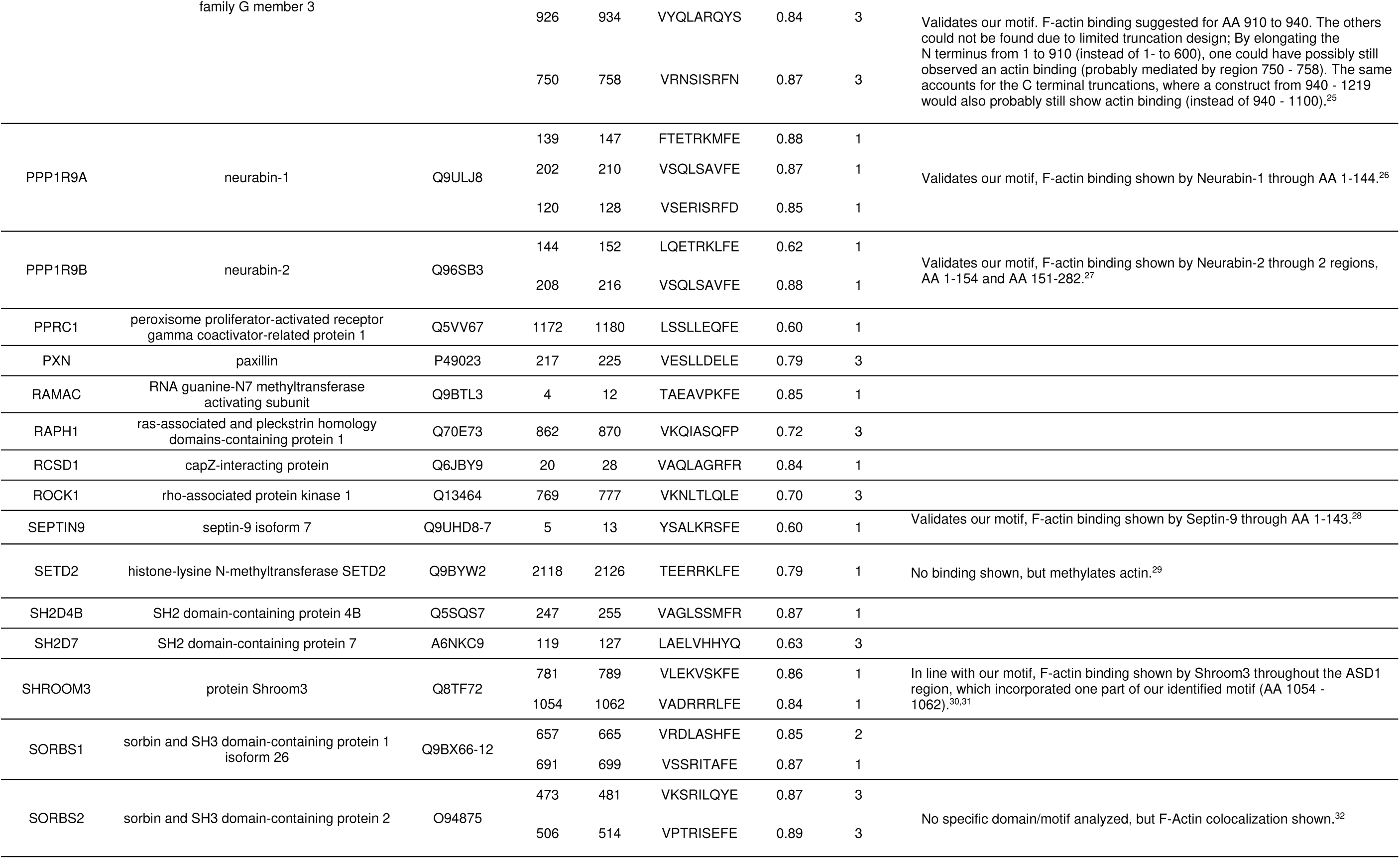
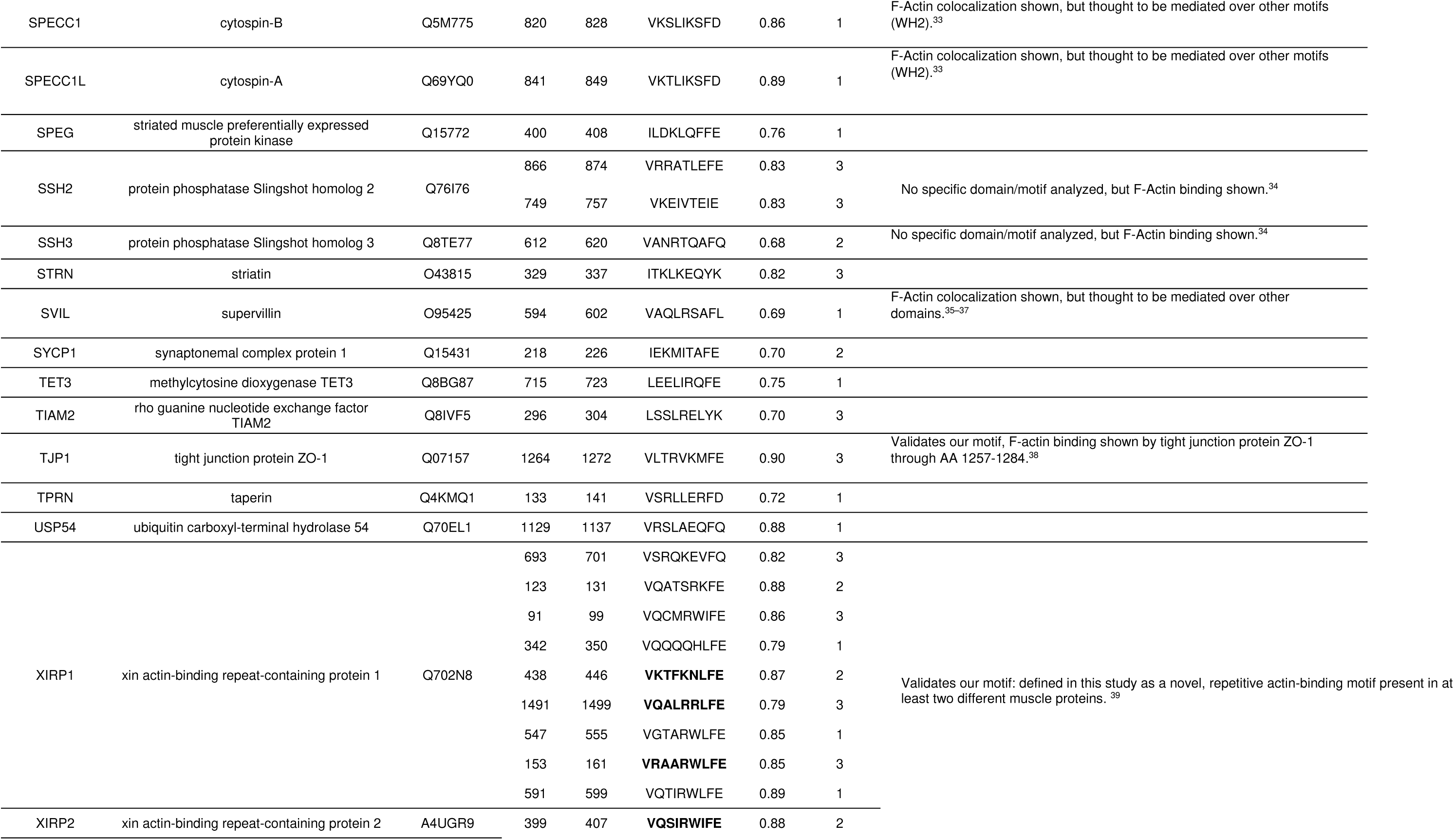
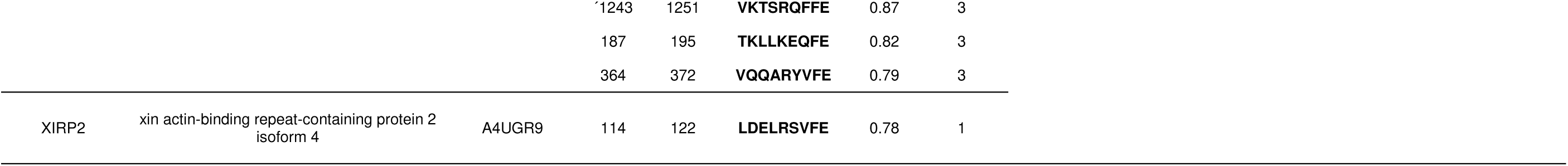
Proteins identified by the SLiMFold pipeline as containing the short linear F-actin–binding motif (SFM). Each entry includes the gene symbol, protein name, UniProt ID, motif boundaries (Start–End), SFM sequence, the mean ipTM (averaged from the top three SLiMFold models), the SLiMFold iteration in which it was identified, and a brief comment providing functional insight or references.

In order to get information about the biological role of the new SFM-proteins, Gene Ontology analysis and manual database research, including Uniprot, NCBI, and PubMed, revealed that they are involved in the regulation of diverse cellular functions (Figure 5b, Extended Data Figure 10 and Table 1). The largest group encompassed the proteins that directly bind to and modulate actin. These include well-described actin modulators, such as Cortactin (CTTN), Caldesmon (CALD1) and Supervillin (SVIL), and many other less investigated proteins. This group is followed by actin regulators, mostly belonging to the RhoGTPase activators (GEFs) or silencers (GAPs). Also, we found microtubule associated proteins, signaling proteins, and proteins involved in the regulation of chromatin, DNA and RNA. The SFM was also identified in proteins, involved in adhesion, junction, focal adhesion, and cilia-proteins.

In conclusion, the SFM is included in highly diverse proteins which is in accordance with our finding that the peptides have arisen spontaneously in evolution.

## Discussion

Actin binding proteins (ABPs) are essential for the control of cellular actin dynamics, and mutation, mis-expression or deletion of ABPs can induce severe diseases^4–8^. The biological role of ABPs is defined by their actin binding mode, and its targeting to actin can alter cytoskeletal dynamics, enzyme activity, or cellular localization (reviewed in ^86,87^). In this study, we present the discovery of a new short linear F-actin binding motif (SFM), and provide first evidence for its biological role.

Detailed mutation analysis revealed that inside the SFM VxxLxxxFE, the valin (P1) and phenylalanine (P8) are essential to bind to F-actin while the amino acids in between modulate affinity, thus are important. This finding was confirmed by evolution analysis. Also, a low dN/dS ratio at these positions evidenced purifying selection and a strong evolutionary drive to retain functional F-actin-binding capacity.

Moreover, phylogenetic analysis indicates that the SFM emerged spontaneously (*ex nihilo*) rather than via exon shuffling or domain duplication. Most likely the motif randomly developed from point mutations, and since it provided a new functionality, it was negatively selected. This *ex nihilo* development seems to be typical for SLiM evolution, among which classical examples are nuclear import and export motifs as well as the calmodulin binding site^88^. Similar to the SFM, they target highly diverse proteins to cellular destinations or mediate binding of protein interaction partners (reviewed in ^89^).

Accordingly, the SFM-containing proteins can be categorized into eight main groups: (1) regulation of microtubule, (2) cilia, (3) cell junctions, (4,5) actin, (6) signal transduction (7) nuclear structure and RNA processing, (8) chromatin regulation and DNA repair. Among these, the SFM-containing proteins involved in actin regulation encompass the highest number of proteins. These include actin binding proteins known to bind to actin via canonical actin binding motifs such as CTTN^52,90^, FHOD3, SVIL, MYO18A, MYO15B and CALD1 as well as RhoGEFs and RhoGAPs whose actin binding activity is partly described.

Since the SFM only mediates weak F-actin binding, we assume that it has a synergistic or additive effect on proteins targeted to actin by canonical actin binding domains. Indeed, for CTTN (cortactin) it has been shown that in addition to the cortactin repeats an adjacent linker, which includes the SFM, is required to mediate high affinity binding to F-actin^52^. Since cortactin stabilizes Arp2/3 F-actin branches, this activity may be important for the maintenance of cellular protrusions. Also, for CALD1 (Caldesmon) the SFM has been identified, and it has been shown that its binding to actin could be displaced by calmodulin, indicating that SFM-mediated actin binding is weakened after cellular stimulation^50^. A synergistic or additive effect of the SFM on actin binding may also work in actin binding proteins containing SFM repeats, including XIRP1, XIRP2, PPP1R9B, and SHROOM3. Also, the actin-regulating Rho-GTPase PLEKHG3 include SFM repeats, and its deletion abrogates cell polarity of 3T3 fibroblasts^61^. Moreover, depletion of the SFM-based actin-binding site in the RhoGEF ARHGEF11 disrupts the balance between Rac and Rho signaling, impairing the formation of leading-edge protrusions and trailing-edge retractions in cancer A431 cells^91^. Likewise, SFM repeats are present in the signaling proteins, SORBS1, SORBS2, and SSH2.

However, ITPKA only includes one SFM, but its deletion had strong effects on the actin architecture as well as on invasion of lung cancer cells^37^, showing that SFM repeats are not essentially necessary for actin targeting. Also, ITPKA is a good example for actin binding proteins involved in the regulation of signal transduction. Deletion of the ITPKA-SFM significantly reduced the InsP_3_Kinase activity of ITPKA^36^, and thereby alters calcium signaling in hippocampal neurons and cancer cells^10,34^. Likewise, the cancer-related protein kinase PEAK is targeted to actin, and phosphorylates the focal adhesion protein paxillin, indicating that its actin targeting coordinates the architecture of focal adhesions after integrin activation^60^. In lymphocytes, HCLS1 is a substrate of tyrosine kinases, and activates the Arp2/3 complex after SFM-controlled actin binding^54^. Thus, it seems that SFM-mediated actin targeting represents a signaling hub integrating actin dynamics and cellular signal transduction.

Moreover, for ZO-1 (TJP1) it could be shown that deletion of its SFM alters the barrier function in kidney MDCK cells, indicating that the SFM exhibits an important function in connecting the actin cytoskeleton with cellular junctions^67^. However, currently there are no studies analyzing actin targeting of SFM-proteins involved in the regulation of cilia, microtubule, chromatin modification, DNA repair, nuclear structure and RNA processing. These groups encompass proteins with broad cellular function such as the kinesins KIF21B and KIF23, as well as the DNA demethylating TET3, and it will be very exciting to elucidate their role in actin targeting in future studies. Finally, two main questions should be addressed in future studies: (1) Does the SFM provide a broad cellular network, or (2) does it define local molecular circuits refined during evolution to adapt to environmental changes? Since the SFM may have developed *ex nihilo,* the second hypothesis is more likely. However, future studies are necessary to prove this assumption.

In conclusion, SFM-mediated F-actin targeting regulates the properties of diverse cellular proteins, and we strongly assume that future studies will uncover yet not identified biological roles. Moreover, expanding our motif search to include non-human databases could reveal the conservation and functional diversity of SFMs across different species, providing deeper evolutionary insights and potentially identifying novel SFM-containing proteins. Looking forward, we envision that the elucidation of the biological role of SFM-proteins will uncover yet unknown cellular regulation mechanisms involved in physiological and pathological settings.

## Methods

### Pipeline Overview

The identification of F-actin-binding SLiMs was performed using the SLiMFold pipeline, which integrates multiple bioinformatic tools to systematically identify, filter, and validate candidate motifs. The pipeline was executed in three sequential iterations, refining results at each step to improve specificity and accuracy.

#### 1. Prerun

The process begins with **Prerun.ipynb**, where hypothesized SLiMs (in this work, the aligned sequences of ITPKA and Lifeact ABDs from P1 to P9) were used along with the BLOSUM62 substitution matrix^92^ to create a position-specific scoring matrix (PSSM). This PSSM was used to scan the human proteome (NCBI Taxonomy ID: 9606) for putative motif hits. To further filter out false positives, we incorporated the following criteria:

- **PSSM Score**: The matrix was used to score sequences in the database, identifying motifs similar to the hypothesized SLiMs^93^.
- **IUPRED**: Predicts intrinsically disordered regions, highlighting regions suitable for motif embedding. The mean IUPRED score was calculated for the motif and its 60-residue flanking regions (N- and C-terminal)^43^.
- **ANCHOR**: Identifies regions in intrinsically disordered regions likely to bind structured partners. The mean ANCHOR value was calculated within the motif^42^.
- **PSIPRED**: Predicts the probability of secondary structures (helix, beta strand, and random coil)^41^.

All cut-offs for these criteria were progressively relaxed across iterations (see below). This approach allowed the identification of additional hits while iteratively strengthening the PSSM, reducing false positives. Each hit was extended by ±20 flanking residues for subsequent homology searches, and redundant sequences were removed. Each hit was then paired with the bait sequence (in this study, the human actin sequence) and saved as a separate FASTA file, formatted for direct compatibility with ColabFold.

For each hit, multiple sequence alignments (MSAs) were generated using **jackhmmer**^44^ with the UniRef90 database. The following jackhmmer parameters were applied: number of iterations: 5, e-value: 1e-5, no f1 or f2 filter (Original AF2 code: number of iterations: 1, e-value: 1e-5, f1: 0.0005. f2: 0.00005). The modified jackhmmer parameters improved the number of sequences retrieved, likely enhancing detection of more distant homologs. Additionally, parallelization was implemented to process multiple hits simultaneously, significantly speeding up the alignment step. The sto-file for the bait sequence was constructed in the same manner. Output alignments were reformatted to A3M format using the **HH-suite’s reformat.pl**^94^ script to ensure compatibility with downstream structure prediction tools. Alignments were sorted and combined to meet AlphaFold2 Multimer input criteria.

#### 2. ColabFold looped

The **ColabFold_looped.ipynb** notebook, originally developed by the Steinegger lab^95^, was modified to enable batch processing and the integration of custom MSAs, making it more suitable for the SLiMFold workflow. These modifications were designed to streamline the process and enhance flexibility when handling multiple input sequences. In the modified version, the notebook was adapted to loop through all FASTA-files in a specified folder, such as one located in Google Drive. This automation allowed for seamless processing of multiple sequences without the need for manual input, significantly improving the workflow’s efficiency. Additionally, the modified notebook incorporated the ability to retrieve precomputed A3M files from another designated folder. These alignments, created during the Prerun.ipynb step, were matched to their corresponding FASTA-files based on their names. This ensured accurate integration of the custom MSAs into the ColabFold predictions. To manage outputs effectively, the notebook allowed users to specify a folder for storing results. This organizational setup made it easier to handle and analyze the results of large-scale computations.

Compared to the original ColabFold batch notebook, these modifications introduced enhanced flexibility by supporting custom MSAs and allowing adjustments to the number of seeds used in the AlphaFold2 predictions.

#### 3. Postanalysis

The post-analysis step was performed using the **Postanalysis.ipynb** script, which systematically processed and analyzed the outputs from the SLiMFold pipeline. This step integrated sequence and structural data to evaluate the conformational and functional relevance of candidate F-actin-binding SLiMs. The pLDDT, pTM, and ipTM scores were extracted to assess prediction confidence and interaction reliability of each predicted structure. Structures with ipTM scores below 0.6 were excluded from further analysis, ensuring a focus on high-confidence predictions. Each remaining structure was aligned to a reference PDB file (in this study, the ITPKA-actin complex) to evaluate conformational similarity.

The **root-mean-square deviation (RMSD)** between the alpha carbon (Cα) atoms of the predicted structure and the reference structure at the motif positions (P1 to P9) was calculated to quantify structural similarity. RMSD was computed as:

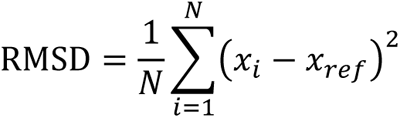

where *N* is the number of aligned residues, *x_i_* represents the atomic coordinates of a residue in the predicted structure, and *x_ref_* represents the corresponding coordinates in the reference structure.

Angular analysis was performed using vector-based calculations, where the spatial coordinates of the alpha carbon (Cα) atoms of key residues in the SLiM (P1 to P9) were used to define vectors representing the orientation of the motif. Two angles, **phi** (φ, azimuthal angle in the x-y plane) and **theta** (θ, polar angle relative to the z-axis), were calculated to quantify the motif’s orientation relative to the reference vector:

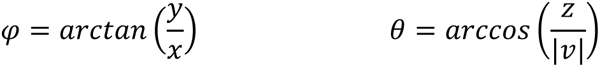

where *x* and *y* are the coordinates of the vector projection in the x-y plane and where *z* is the z-coordinate of the vector and |ν| is the magnitude of the vector. Helix polarity was also calculated to capture the directionality of the predicted motif’s structural alignment. The **delta phi (Δ_φ_)** and **delta theta (Δ_θ_)** values, representing the differences between the angles of the predicted and reference structures, were calculated for each candidate SLiM. These metrics, along with RMSD and polarity, provided a comprehensive structural profile for clustering.

Clustering was performed using the HDBSCAN^45^ algorithm, leveraging RMSD, **Δ_φ_**, **Δ_θ_**, and polarity as input features. The clustering parameters, including minimum cluster size and minimum samples, were optimized using the following metrics:

- **Silhouette Score**: Evaluates cluster separation and cohesion.
- **Davies-Bouldin Index**: Measures intra-cluster similarity relative to inter-cluster separation.
- **Calinski-Harabasz Index**: Assesses the ratio of between-cluster dispersion to within-cluster dispersion.

Optimal clustering parameters were selected based on these indices, ensuring robust and meaningful grouping of structurally similar SLiMs. For further analysis, all structures belonging to a single cluster were exported as a PyMOL session file (.pml), enabling manual inspection and visualization. For each cluster, the corresponding sequences were compiled into a FASTA file. These sequences were used to generate sequence logos^96^, highlighting conserved residues and identifying potential functional motifs. Additionally, each sequence was mapped to its corresponding gene using the NCBI database. This mapping facilitated gene ontology (GO) enrichment analysis, allowing functional insights into the biological roles of the identified SLiMs.

### Cloning strategies

All constructs were generated via PCR-based cloning using high-fidelity polymerases (Q5 or Phusion; M0491S or M0530S, New England Biolabs Inc., Massachusetts, USA) following the manufacturer’s instructions. For inserting longer DNA fragments such as full-length protein sequences, we employed T4 SLIC^97^ (Sequence and Ligation Independent Cloning) (T4 DNA Polymerase; M0203S, New England Biolabs Inc., Massachusetts, USA), which relies on ∼30 nucleotide overlaps between vector and insert for seamless assembly. Short peptides or specific mutations were introduced using a “QuickChange-like” site-directed mutagenesis protocol, wherein primer pairs incorporated short (8–12 nt) overlaps at their 5′ ends; the PCR products were subsequently treated with DpnI (R0176S; New England Biolabs Inc., Massachusetts, USA) to remove the template and then transformed into *E. coli* XL1-Blue (200249, Agilent Technologies, Inc., California, USA). Alternatively, the PCR products were processed using the KLD (Kinase–Ligase–DpnI) enzyme mix (M0554S, New England Biolabs Inc., Massachusetts, USA) to remove the template and ligate the new amplicon. For bacterial expression, PCR products were cloned into pSF421-based expression vectors (e.g., pSF421_10xHis_GFP_TEV), while for eukaryotic expression in mammalian cells (e.g., H1299), target genes were inserted into an mEGFP-N1 backbone. All final plasmids were verified by Sanger sequencing (Microsynth Seqlab GmbH, Göttingen, Germany). Verified constructs were subsequently transformed into *E. coli* Rosetta™ (DE3) (Novagen, Merck KGaA, Darmstadt, Germany) for bacterial protein expression or transfected into mammalian cells. Complete plasmid lists, vector maps, sequencing results, and primer sequences are provided in Supplementary Tables 2 and 3 and Source Data.

### Purification of Actin

Actin was prepared from Gallus gallus (chicken) skeletal muscle as described in reference^98^. The final purification step was performed using a HiLoad 16/600 Superdex 200 column equilibrated with G-actin buffer (5 mM Tris-HCl, pH 7.5, 0.2 mM CaCl_2_, 0.5 mM DTT and 0.2 mM ATP).

### Expression and Purification of ITPKA and GFP-Peptides

All constructs were expressed in *E. coli* Rosetta™ (DE3) competent cells (Novagen, Merck KGaA, Darmstadt, Germany). Cells were grown in Terrific Broth (TB) medium at 37°C to an optical density (OD_600_) between 0.6 and 1.0. Protein expression was induced with 0.2 mM isopropyl β-D-1-thiogalactopyranoside (IPTG) at 18°C for 16 hours for ITPKA and with 0.2 mM isopropyl β-D-1-thiogalactopyranoside (IPTG) at 37°C for 4 hours for the GFP-Peptides. After induction, cells were harvested by centrifugation at 3,000 × g for 20 minutes at 4°C. The cell pellets were resuspended in cold phosphate-buffered saline (PBS) buffer, centrifuged at 2,000 × g for 30 minutes at 4°C, flash-frozen in liquid nitrogen, and stored at −80°C.

For protein purification, the cell pellets were resuspended in ice-cold Buffer A (50 mM Tris-HCl, pH 7.4, 400 mM NaCl, 3 mM MgCl_2_ 1 mM β-mercaptoethanol for ITPKA and 50 mM Tris-HCl, pH 7.4, 300 mM NaCl, 1 mM β-mercaptoethanol for GFP-Peptides) and homogenized using an IKA® ULTRA-TURRAX® disperser (IKA-Werke GmbH & Co. KG, Staufen, Germany). DNase I was added, and the cells were lysed with a Constant Cell Disruption System (Constant Systems Limited, Daventry, UK) at 1.8 kbar. Following lysis, phenylmethylsulfonyl fluoride (PMSF) was added to a final concentration of 1 mM and imidazole to a final concentration of 25 mM. Cell debris was removed by centrifugation at 43,000 × g for 30 minutes at 4°C, and the supernatant was incubated with Ni-NTA agarose resin (SERVA Electrophoresis GmbH, Heidelberg, Germany) for 30 minutes at 4°C. The bound proteins were eluted using a gradient of Buffer B (Buffer A with 500 mM imidazole). Fractions were analyzed by SDS-PAGE, and those with >90% purity were pooled and then dialyzed overnight at 4°C in Buffer A.

For the ITPKA protein, an additional cleavage step was performed to remove His tag. The pooled protein fractions were dialyzed overnight at 4°C in the presence of TEV protease, at a ratio of 1 mg TEV protease per 40 mg protein substrate, in Buffer A. Following dialysis, the cleaved protein was separated from the tags and TEV protease by reverse immobilized metal affinity chromatography (IMAC) using a Ni^2+^-NTA column (Bio-Rad Laboratories, California, USA) on an NGC™ Liquid Chromatography System (Bio-Rad Laboratories, California, USA).

Subsequently, the pooled reversed IMAC fractions (for ITPKA) or the dialyzed proteins (for GFP Peptides) were concentrated to 0.5–2 mL using Amicon® Ultra Centrifugal Filters (Merck-Millipore, Massachusetts, USA) and further purified by size-exclusion chromatography using a HiLoad Superdex 16/600 200pg Gel Filtration Column (Cytiva, Massachusetts, USA) on an NGC™ Liquid Chromatography System (Bio-Rad Laboratories, California, USA), equilibrated with Buffer D (50 mM Tris-HCl, pH 7.4, 150 mM NaCl, 3 mM MgCl_2_ 1 mM β-mercaptoethanol for ITPKA and 50 mM Tris-HCl, pH 7.4, 150 mM NaCl, 1 mM β-mercaptoethanol for GFP-Peptides). The peak fractions were concentrated, aliquoted, then flash-frozen in liquid nitrogen, and stored at −80°C.

### Cryo-EM data acquisition and processing

We initiated all sample preparations using frozen aliquots of G-actin. To convert G-actin into F-actin, we first thawed the samples and performed centrifugation at 100,000 x g for 30 minutes to eliminate potential aggregates. The polymerization process involved adding F buffer to achieve a final concentration of 20 mM Tris-HCl pH 7.5, 100 mM KCl, 2 mM MgCl_2_, 0.5 mM ATP, 1 mM EGTA, and 1 mM DTT, followed by incubation for one hour at room temperature. Subsequently, we collected the filaments by centrifuging at 100,000 x g for one hour at 4°C and resuspended them in F buffer supplemented with Phalloidin at twice the molar concentration of F-actin.

For cryo-EM grid preparation, we used Quantifoil 200-mesh 2.0/1.0 holey carbon grids. Specifically, for Phalloidin-stabilized F-actin, we applied 3.5 μl of 1 μM solution onto glow-discharged grids. For the Phalloidin-stabilized F-actin-ITPKA complex, we initially applied 2 μl of 1 μM Phalloidin-stabilized F-actin to the grids. Subsequently, we added 2 μl of 10 μM ITPKA and mixed it directly on the grids immediately before plunge-freezing in a liquid ethane/propane mixture using a Vitrobot Mark IV (FEI). Datasets were collected on an FEI Titan Krios (300 KV; Gatan K3 camera; pixel size: 0.826 Å; dose per frame:1.4; defocus range: 0.8 to 3.0 μm).

Single-particle helical reconstruction was performed using Relion 4^99^. Movies were firstly motion-corrected using MotionCor2^100^, and then the contrast transfer function (CTF) estimation was done using CTFFIND4^101^. Particle segments picking model was trained in cryolo 1.7^102^. The helical segments were extracted into 360 × 360 boxes and the junk segments were excluded after 2D classification. The initial helical parameters of a helical rise of 27.3 Å and a helical twist of −166.5° were applied for 3D classification and autorefinement in Relion 4.0. Overall, gold-standard resolution (Fourier shell correlation = 0.143) was calculated in Relion 4.0. The statistics for data collection and processing are listed in Supplementary Table 4.

### Model building and refinement

Previously published models of phalloidin bound F-actin (PDB: 6T20^103^, 7BTI^40^), and the AlphaFold2 structure of ITPKA were used as initial models. Models were initially built in ChimeraX 1.7^104^, and further refined against the cryo-EM maps using ISOLDE^105^, and real space refinement in Phenix^106^. The detailed model information and validation statistics for final models are described in Supplementary Table 4.

### Cell culture

NCI-H1299 (H1299) cells were kindly provided by Cagatay Günes (Hamburg, Germany). For detailed cellular characteristics, refer to the American Type Culture Collection (ATCC, Rockville, USA). The cells were cultured in Dulbecco’s Modified Eagle’s Medium (DMEM) supplemented with 10% (v/v) fetal calf serum (FCS), 4 mM L-glutamine, 100 μg/ml streptomycin, and 100 U/ml penicillin. The culturing conditions were rigorously maintained to ensure optimal growth and viability. Additionally, the cells were tested regularly for mycoplasma contamination to ensure experiments were conducted with mycoplasma-free cells.

Primary human monocytes were isolated from buffy coats (kindly provided by Frank Bentzien, Transfusion Medicine, UKE, Hamburg, Germany). 20 ml blood was coated on 15 ml Lymphocyte Separation Medium 1077 (PromoCell, Heidelberg, Germany) and centrifuged for 30 min at 4°C and 460×g. Buffy coats were transferred to a new 50 ml Falcon tube and filled up to 50 ml with cold RPMI (Gibco, Paisley, UK). Leukocyte fractions were washed twice in RPMI and centrifuged for 10 min, as described above. Enriched leukocytes were resuspended in 400 μl monocyte buffer (5 mM EDTA and 0.5% human serum albumin in Dulbecco’s PBS [DPBS], pH 7.4), mixed with 100 μl of magnetic beads suspension coupled to anti-CD14 antibodies (Miltenyi Biotec, Bergisch Gladbach, Germany) and incubated for 15 min on ice. The mixture was subsequently loaded onto Separation columns LS (Miltenyi Biotec, Bergisch Gladbach, Germany) that were previously placed in a magnetic holder and equilibrated with 500 μl cold monocyte buffer. Trapped CD14+ monocytes were washed on column with 500 μl monocyte buffer and then eluted with 1ml monocyte buffer into 15 ml cold RPMI after removal from the magnets. After centrifugation for 10 min at 4°C and 460×g, the supernatant was removed and cells were resuspended in 40 ml RPMI and seeded on a 6-well plate (Sarstedt, Nuembrech, Germany) at a density of 2*106 cells per well. After adhesion of monocytes for 1h, RPMI medium was replaced by 2 ml monocyte culture medium (RPMI substituted with 20% human serum and 1% penicillin/streptomycin (Sigma-Aldrich, Missouri, USA). Monocytes were cultivated in an incubator at 37°C, 5% CO2, and 90% humidity. Isolated monocytes were differentiated for at least 6 days.

### Transfection of Cells

For H1299 cells, a total of 2.5 × 10^4^ cells were seeded into 8 well chamber slides (Ibidi, Gräfelfing, Germany). After 16 h, the cells were transfected with 0.5 µg with any mEGFP-N1-Peptide or mEGFP-N1-FL-Proteins using the K2® Transfection Reagent (T060-0.75, Biontex Laboratories GmbH, München, Germany) according to the manufacturer’s instructions. After 24 h, the cells were fixed with 4% paraformaldehyde/4% sucrose, stained with rhodamine-conjugated phalloidin (ab235138, Abcam Limited., Cambridge, UK), and analyzed by fluorescence microscopy using the Olympus IXplore Live microscope imaging system and the FV3000 confocal laser scanning microscope (Olympus Corporation, Tokyo, Japan).

Macrophages were detached by incubation of Accutase (Invitrogen, Massachusetts, USA) for at least 30 min in culturing conditions. Cells were collected with monocyte culture medium, washed in PBS pH 7.3 and resuspended in R-Buffer (at a concentration of 106 cells per 100 µl buffer, 10 µg DNA), provided by the Neon Transfection System (Invitrogen, Massachusetts, USA). Macrophages were transiently transfected with plasmid DNA with the following settings: voltage 1,000 V; width 40 ms, and 2 pulses. Transfected cells were resuspended in RPMI and seeded on 12-mm glass coverslips (105 cells per coverslip). The cells were left to adhere for 1 h at culturing conditions. Thereafter, 1 ml of monocyte culture medium was added, and cells were incubated overnight.

### Immunofluorescence and microscopy

Cells were fixed for 10 min in 3.7% formaldehyde in PBS, and permeabilized for 10 min in PBS containing 0.5% TritonX-100. Thereafter cells were incubated for 60 min in blocking solution (2% BSA in PBS) with 1:400 phalloidin-568. Cells were washed three times in PBS and mounted on glass slides with FluoromountG (Invitrogen Massachusetts, USA) containing DAPI (Sigma-Aldrich, Missouri, USA). Images of fixed samples were acquired with a confocal laser-scanning microscope Olympus FV3000 equipped with an 60x UPlanApo HR Oil objective and Olympus FV-3000 software.

### Poji Macro Analysis

Semi-automated Poji analysis was performed as previously described in reference^107^. Profile analysis was performed on ROI of transfected cells with podosomes defined by phalloidin-568 staining with a circle size of single isolated podosomes of 25 pixels. For more detailed information, see https://github.com/roherzog/Poji.

### Cosedimentation assay and *K_D_* Determination

The dissociation constant (*K_D_*) and binding strength of the interaction between the actin-binding protein (ABP) and filamentous actin (F-actin) were determined using an F-actin cosedimentation assay.

For this, preparations were made using frozen aliquots of G-actin. To convert G-actin into F-actin, the samples were thawed and centrifuged at 100,000 × g for 30 minutes to eliminate potential aggregates. The polymerization process involved adding F-buffer to achieve a final concentration of 20 mM Tris (pH 7.5), 100 mM KCl, 2 mM MgCl_2_, 0.5 mM ATP, 1 mM EGTA, and 1 mM DTT. The samples were then incubated for one hour at room temperature. The resulting F-actin was mixed with an equivalent amount of ABP and incubated for one hour at room temperature before cosedimentation at 100,000 × g for 30 minutes. Pellet and supernatant fractions were resuspended in sample buffer, heated at 95 °C for 15 min, and analyzed by SDS-PAGE and Western blot or Coomassie staining as detailed below.

For ABPs exhibiting relatively **low affinity** (*K_D_* > [*ABP*], such that free actin remains largely undepleted), we measured only the pellet fraction by Western blot and plotted densitometry (D) as a function of the total actin concentration [*Actin*]_total_. These data were fit in GraphPad Prism using the “One site – Specific binding” hyperbolic model:

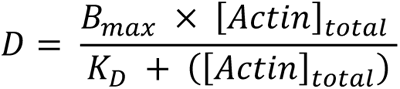

Here, *B_max_* corresponds to the maximal densitometry signal at saturation.

For **higher-affinity** ABPs (*K_D_* < [*ABP*]), the free actin can be substantially depleted, so a simple hyperbolic fit may be inaccurate. In these cases, we quantified both pellet and supernatant fractions by SDS-PAGE to determine the fraction of total ABP bound, *Y*. We then fit *Y* against [*Actin*]_total_. using the „Quadratic” (or “Tight Binding”) model^108^:

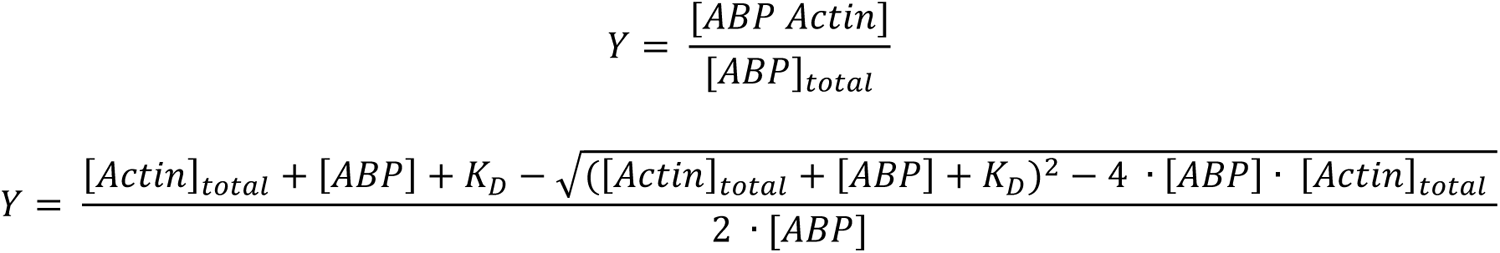

All experiments were performed in triplicate to ensure reproducibility. Varying concentrations of actin (0–120 µM) were incubated with a fixed concentration of ABP (4 µM). In each replicate, a negative control (0 µM actin) verified background signal. For low-affinity peptides, blot exposures were selected to remain in the linear range of detection, and densitometry signals (D) were directly used in the hyperbolic fit. For high-affinity assays, we confirmed mass balance by checking that pellet + supernatant matched the total ABP added. Nonlinear regression in GraphPad Prism was used to extract *K_D_* and *B_max_*.

For the binding strength determination of USP54 and SHROOM3 mutants (M1, M2, M3, and M4), the proteins were employed at a final concentration of 4 µM with F-actin at 12 µM. Densitometric analysis was performed on the Western blot results, quantifying the signal intensity of the bound ABP in the pellet fraction to assess differences in binding strength among the mutants. The densitometry values were corrected by subtraction of the negative control (SFM with 0 µM Actin).

### SDS-PAGE and Western Blotting

Equal volumes of samples were loaded onto 12% polyacrylamide gels for SDS-PAGE. Electrophoresis was performed at 120 V until the dye front reached the bottom of the gel. SDS-PAGE was transferred onto nitrocellulose membranes using a wet transfer system at 60 V for 90 minutes in 1× blot buffer. Membranes were blocked with 5% (w/v) non-fat dry milk or 5% BSA in Tris-buffered saline with Tween 20 (TBST) for 1 hour at room temperature to block nonspecific binding sites. The membranes were incubated overnight at 4°C with the primary antibody specific to GFP (mouse anti-GFP, 1:1,500, Article No. 11814460001, Roche Applied Science, Penzberg, Germany) or actin (rabbit anti-actin, 1:10,000, Article No. A2066, Sigma Aldrich, St. Louis, MO, USA) diluted in blocking buffer. After three washes with TBST, the membranes were incubated for 1 hour at room temperature with the appropriate HRP-conjugated secondary antibody. Goat anti-mouse secondary antibody was used for GFP detection (1:10,000 dilution in TBST), while goat anti-rabbit secondary antibody was used for actin detection (1:10,000). Protein bands were detected using a chemiluminescence reagent (Cytiva Amersham™ ECL™ Prime Western Blot Detection Reagent) and imaged using the INTAS ECL CHEMOCAM (INTAS Science Imaging Instruments GmbH, Göttingen, Germany). Band intensities were analyzed using ImageJ^109^.

### Position specific frequency matrix (PSFM)

We developed a pipeline for generating position-specific frequency matrices (PSFMs) that quantify amino acid conservation at each position across a set of sequences, focusing specifically on peptides with high affinity or strong colocalization. Multiple sequence alignments (MSAs) derived from SLiMFold’s jackhmmer output were manually curated to remove alignments with names indicating different proteins, ensuring retention of only true homologs. These curated MSAs served as input for generating individual frequency tables for each peptide, which were then averaged to account for variations in the number of homologous sequences. A 23×N matrix (where N is the sequence length) was generated, with each element representing the proportion of a specific amino acid at a given position. To reduce skewness in the data and emphasize subtle conservation trends, a square root transformation was applied to the matrix. Heatmaps of the PSFMs were visualized using matplotlib, with conserved positions annotated and residue indices labeled for interpretation. Detailed code is provided in the reference^110^.

### Phylogenetic and Motif Analysis

To investigate the evolutionary history of F-actin binding motif-containing proteins and their isoforms, we conducted a comprehensive phylogenetic analysis. Orthologous sequences were retrieved from OrthoDB^111^ and aligned using DECIPHER^112^ (100 iterations, 200 refinements). Maximum likelihood phylogenetic trees were constructed with IQ-TREE 2^113^ (WAG+G model, 1000 ultrafast bootstraps). Isoform sequences were identified and filtered based on header annotations, and their phylogenetic clustering was visualized. Taxonomic information was incorporated to determine the MRCA of isoform clusters. F-actin binding motifs were extracted using regular expressions and filtered based on predicted disorder (IUPRED^43^), anchor regions (ANCHOR^42^), and random coil propensity (PSIPRED^41^). Position-Specific Scoring Matrices (PSSMs) were generated and iteratively used to refine motif identification. The MRCA of motif-containing sequences was determined using isoform-specific trees. Motif distribution was validated by mapping motifs to a taxon-based tree. Motif development across isoforms and taxonomic classes was analyzed using frequency matrices. Sequence logos^96^ were generated from PSSMs. For nucleotide-level analyses, corresponding DNA sequences were retrieved from NCBI, filtered for ambiguous bases and stop codons, and codon-aligned using PRANK^114^. Phylogenetic trees were inferred with IQ-TREE 2^113^ (GTR+G model), and dN/dS analysis was performed using HyPhy^115^. Detailed code and parameter settings are provided in the Source Data.

### GO Enrichment analysis

To elucidate the biological functions and cellular components associated with Short Linear F-actin Binding Motif (SFM)-containing proteins, we performed Gene Ontology (GO) enrichment analysis using ShinyGO^116^. The analysis aimed to identify overrepresented GO terms within three primary categories: Biological Process (BP), Cellular Component (CC), and Molecular Function (MF). We compiled a comprehensive list of SFM-containing proteins identified in this study as the query set. Utilizing ShinyGO with its standard parameters, we input the query set to perform the enrichment analysis. The analysis employed the default settings for multiple testing correction, including the False Discovery Rate (FDR), ensuring that only GO terms with an adjusted p-value below 0.05 were considered significantly enriched.

### AlphaMissense

For each of the 12 proteins examined in this study, the AlphaMissense pathogenicity score^47^ was retrieved using the canonical UniProt ID. The relevant amino acid positions were then manually aligned based on the SFM motif coordinates. Residues flanking the motif (positions −30 to −20 and +30 to +40) served as control values for comparative analysis. After extracting the AlphaMissense scores for all 12 proteins, the mean score at each aligned position was calculated and the standard error of the mean (SEM) determined. Statistical evaluation of differences between motif residues and the flanking control positions was performed via one-way ANOVA with multiple comparisons.

### Data availability

Coordinates and cryo-EM maps for the F-actin structure have been deposited in the Electron Microscopy Data Bank (EMDB) under accession code EMD-18866, with corresponding Protein Data Bank (PDB) entry 8R3H. The F-actin–ITPKA complex has been deposited in the EMDB under accession code EMD-18868 and in the PDB under accession code 8R3J. Additionally, PDB entries 6T20, 7BTI, and 7AD9 were used for structural comparisons. The SLiMFold pipeline code is available at https://github.com/thp42/SLiMFold, and the PSFM code is provided at https://github.com/thp42/PSFM. Source data underlying this study are provided with the paper.

## Acknowledgements

We thank Christine Blechner and Andrea Mordhorst for excellent technical assistance. We also acknowledge Christian Dahlstroem (AG Windhorst, Universitätsklinikum Hamburg-Eppendorf) Nicholas Kley and Joel Fauser (AG Itzen, Universitätsklinikum Hamburg-Eppendorf) for continuous support. We are grateful to Frank Bentzien (Transfusion Medicine, Universitätsklinikum Hamburg-Eppendorf) for providing buffy coats, and to Robert Herzog and Pasquale Cervero for help in validating podosome detection. We thank Malik Alawi, Philipp Dirksen and Susanne Dobler for critical evaluation of our phylogenetic analysis. We further appreciate the support of the UKE Microscopy Imaging Facility (UMIF, DFG code RI_00489 and INST 152/933-1) for advanced microscopy access and image analysis, as well as Martin Aepfelbacher and Aymelt Itzen for continuous support. This project was funded by Erich und Gertrud Roggenbuck-Stiftung (to S.W. and T.P.). It is part of the doctoral thesis of K.S. and was additionally supported by the Deutsche Forschungsgemeinschaft (RTG2771/P4, to S.L.).

