## Extended Data for "Discovery of a new evolutionarily conserved short linear F-actin binding motif" (from another version of this preprint: version 1, 2025-04-16)

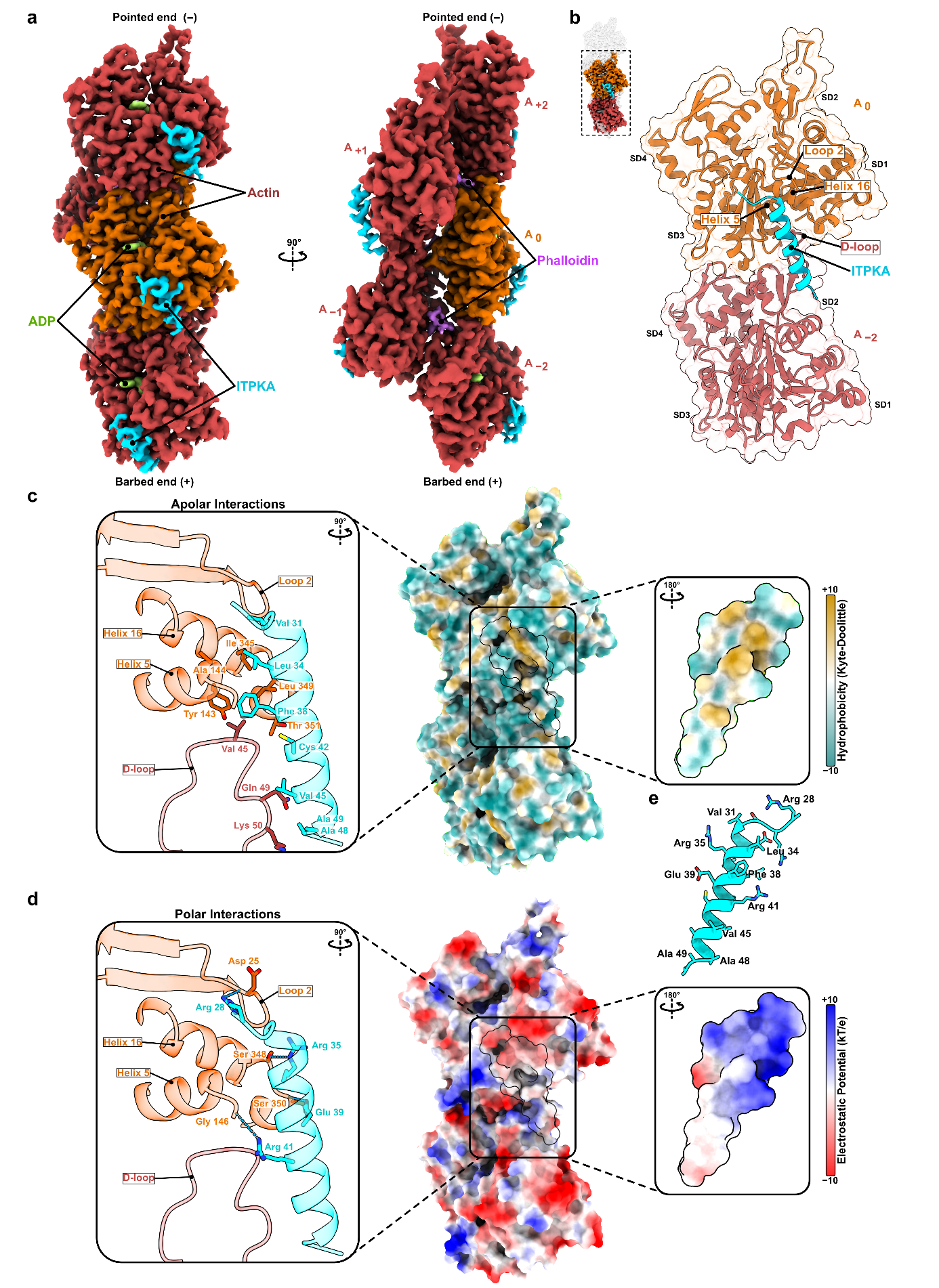
**Extended Data Figure 1: Cryo-EM structure of the ITPKA-F-actin complex.** **(a)** The electron density map illustrates the ITPKA–F-actin complex. A single ITPKA helix of residue 28–49 (cyan) is resolved, forming the primary binding interface with actin. Actin subunits are displayed with the central subunit in orange and surrounding subunits in red. Phalloidin is shown in purple, and ADP in light. **(b)** A cartoon representation of the ITPKA–F-actin complex shows ITPKA bridging two adjacent actin subunits. Key interaction regions in actin helix 5 (AA 137 – 145), helix 16 (AA 338 – 348), loop 2 (AA 21 – 28), the D-loop (AA 38 -51) and actin subdomains SD1–SD4 are labeled. **(c)** Apolar interactions between ITPKA and actin are depicted by surface color-coding for hydrophobicity (yellow = hydrophobic). Left panel: Zoomed-in view and a 90° rotation highlight interacting residues displayed as sticks. Right panel: Zoomed-in view and a 180° rotation; here hydrophobicity scores are shown (low: cyan, high: yellow). **(d)** Polar contacts between ITPKA and actin are shown via electrostatic potential mapping on actin (negative: red, positive: blue). Left panel: Zoomed-in view and a 90° rotation highlight interacting residues displayed as sticks. Right panel: Zoomed-in view and a 180° rotation, here electrostatic scores are shown (negative: red, positive: blue). **(e)** The binding helix of ITPKA is separately depicted as a cartoon with side chains shown as sticks.

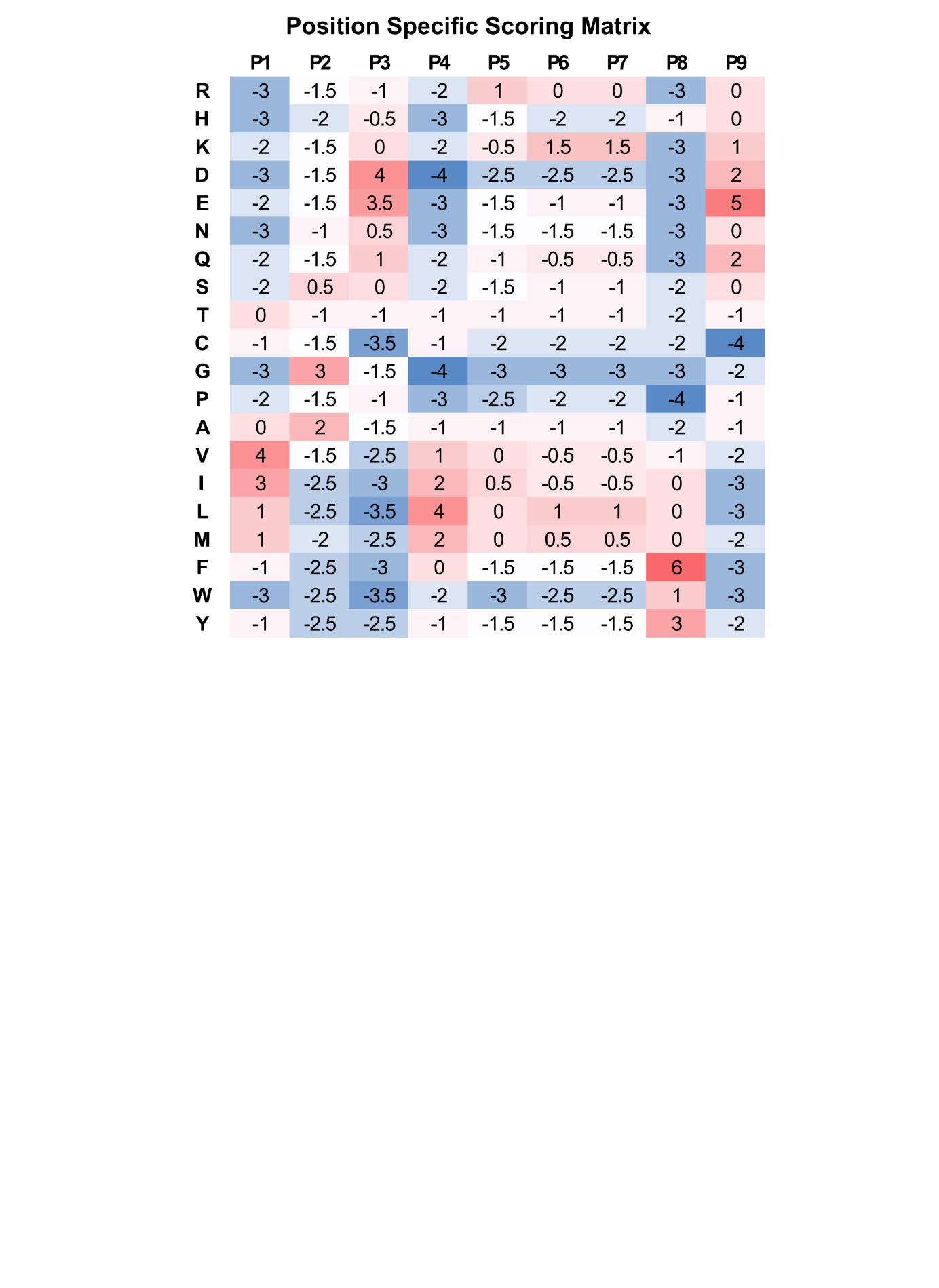
 **Extended Data Figure 2: Position‐specific scoring matrix (PSSM) derived from the two validated motif sequences, ITPKA and Lifeact.** Nine positions (P1–P9) were aligned (see Figure 2c) and scored using a BLOSUM62‐based approach, generating the PSSM shown. In this heatmap, rows are grouped by amino acid type: positively charged (+), negatively charged (-), polar (P), unique (U), non-polar (NP) and aromatic (A) residues and the position‐wise scores are color‐coded (red = favorable, blue = unfavorable). Higher (positive) scores indicate preferential substitutions, whereas lower (negative) values suggest disfavored residues at a given position. This matrix, which reflects sequence conservation in the ITPKA and Lifeact short linear motifs, provided the starting point for screening and iterative refinement in the SLiMFold pipeline (see Figure 2 and Methods).

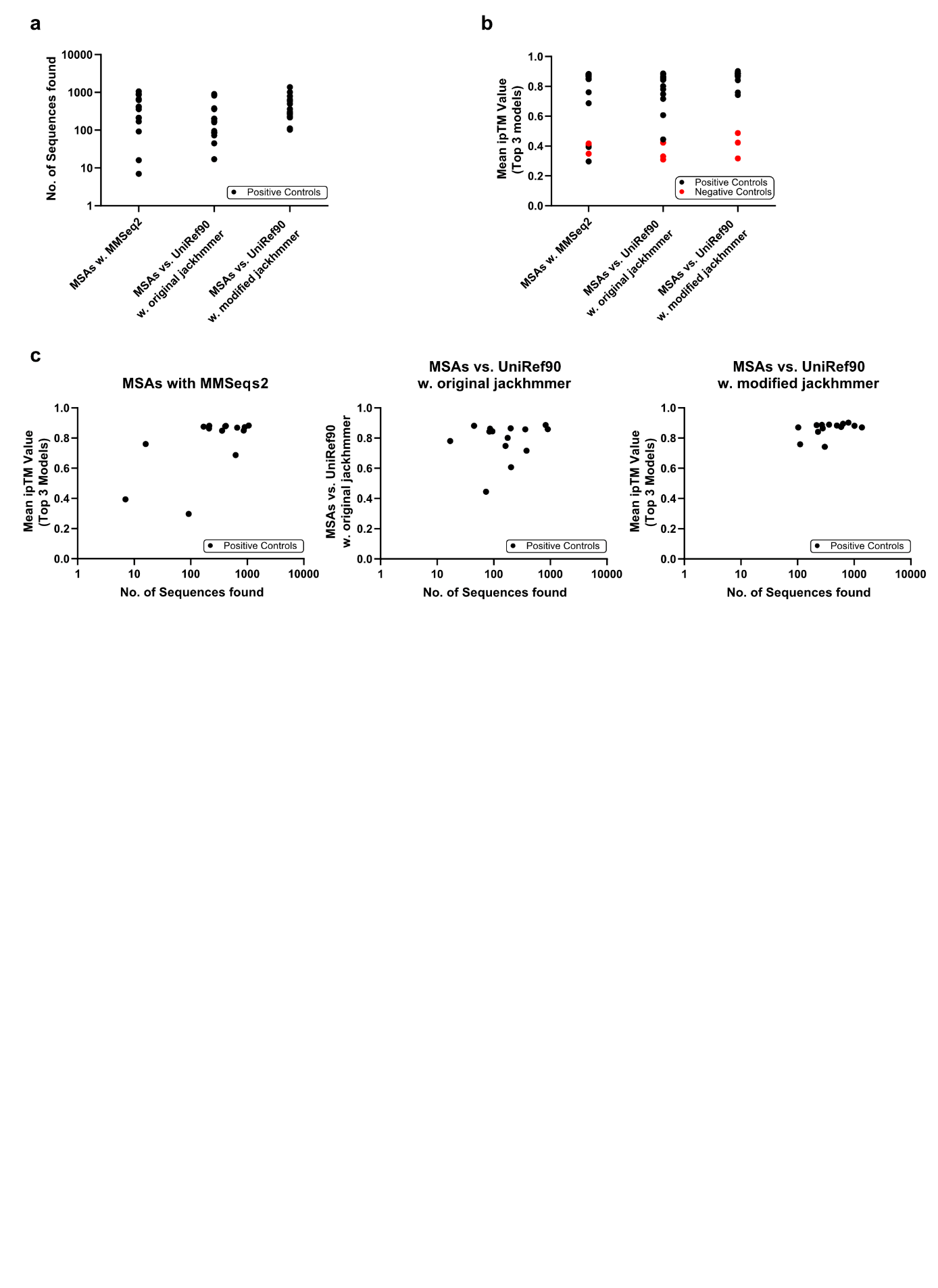
 **Extended Data Figure 3: Optimization of AlphaFold2 parameters for the SLiMFold pipeline. (a)** Comparison of the number of sequences identified (y-axis, log scale) under different multiple sequence alignment (MSA) algorithms: MMSeq2, UniRef90 with original jackhmmer parameters, and UniRef90 with modified jackhmmer parameters. Positive controls (black) are known SLiM-containing peptides validated in this work; negative controls (red) are random amino acid sequences. **(b)** Mean ipTM scores (top 3 ranked of 20 predicted structures) for positive controls (black) versus negative controls (red) across the same MSA methods. Positive controls run with the modified jackhmmer settings exhibit higher mean ipTM values compared to MMSeq2 or the original jackhmmer, suggesting improved interface prediction accuracy with the modified parameters. **(c)** Scatter plots show the relationship between the number of sequences identified (x-axis, log scale) and the mean ipTM values (y-axis) for each MSA approach.

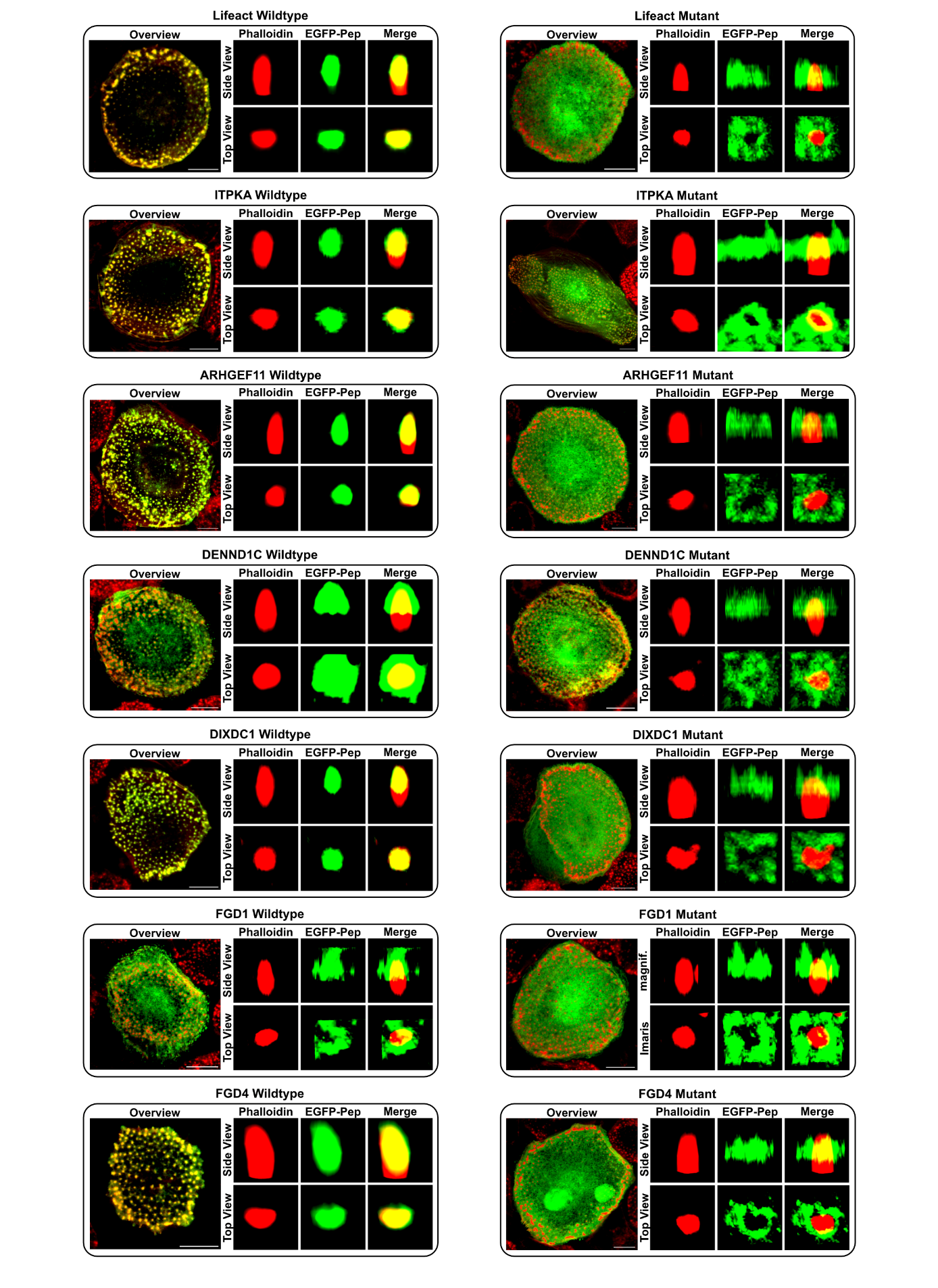
**Extended Data Figure 4: Colocalization of phalloidin-stained actin and EGFP-tagged peptides in macrophage podosomes.** Primary human macrophages were transfected with EGFP‐tagged peptides (green) and stained with phalloidin‐568 (red) to visualize F‐actin at podosomes. Representative confocal images are shown in both top and side views (Imaris 3D reconstructions), emphasizing subcellular peptide localization. Wild‐type peptides generally display robust colocalization with F‐actin (appearing yellow in merged panels), whereas alanine substitutions at key residues (P1 or P8) diminish or abolish F‐actin association. Scale bars, 10 µm.

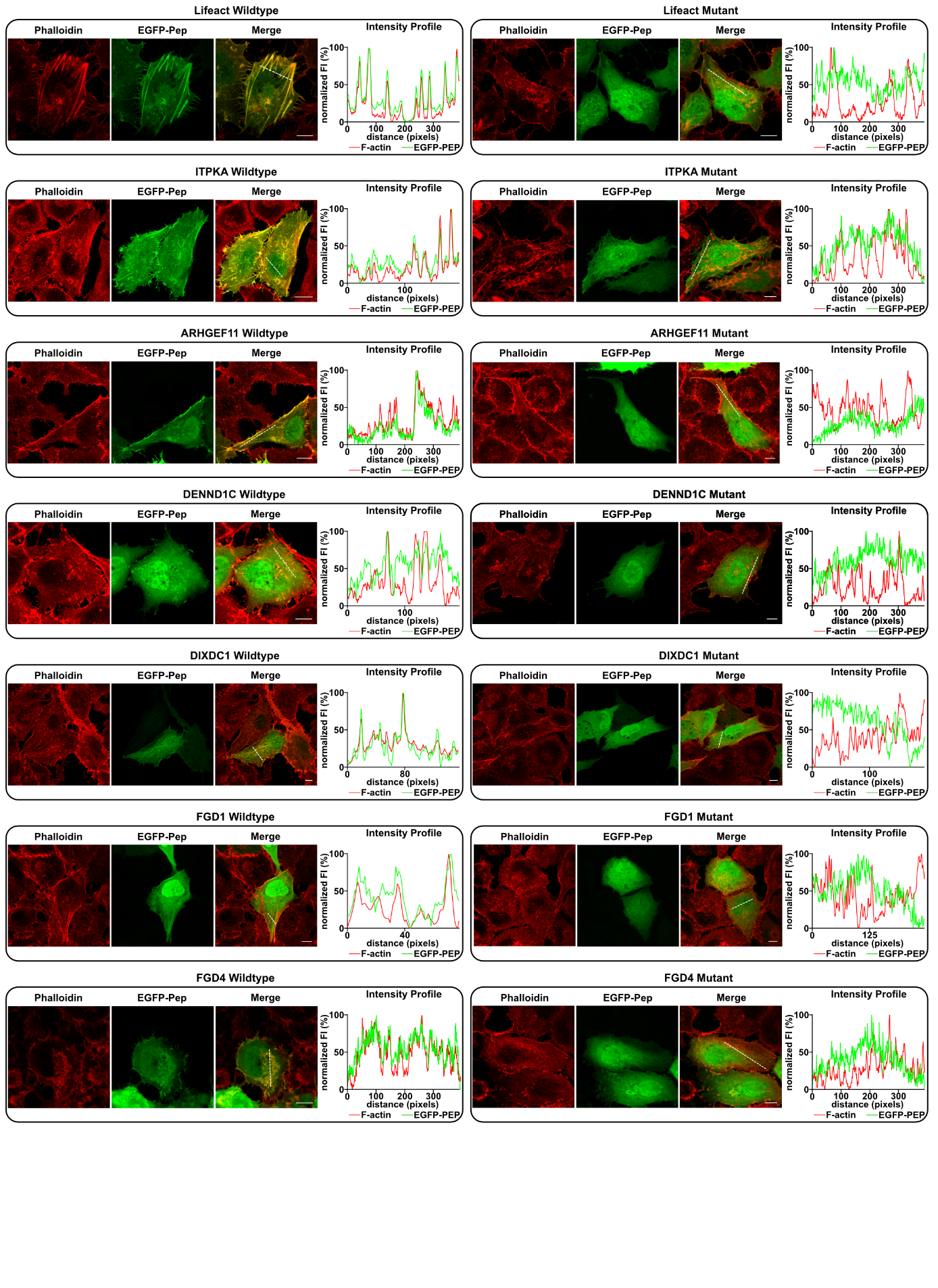
**Extended Data Figure 5: Colocalization of phalloidin-stained actin and EGFP-tagged peptides in H1299 cells , with accompanying intensity profiles.** Representative confocal micrographs show EGFP‐tagged peptides (green) alongside phalloidin‐568 staining of F‐actin (red). Merged images highlight varying degrees of colocalization (appearing yellow). Intensity profiles (right panels) were generated along the dashed white line in each cell, quantifying the overlap between F‐actin and peptide signals. Wild‐type peptides generally display robust colocalization, whereas alanine substitutions at key positions (P1 or P8) diminish or abrogate actin association. Scale bars, 10 µm.

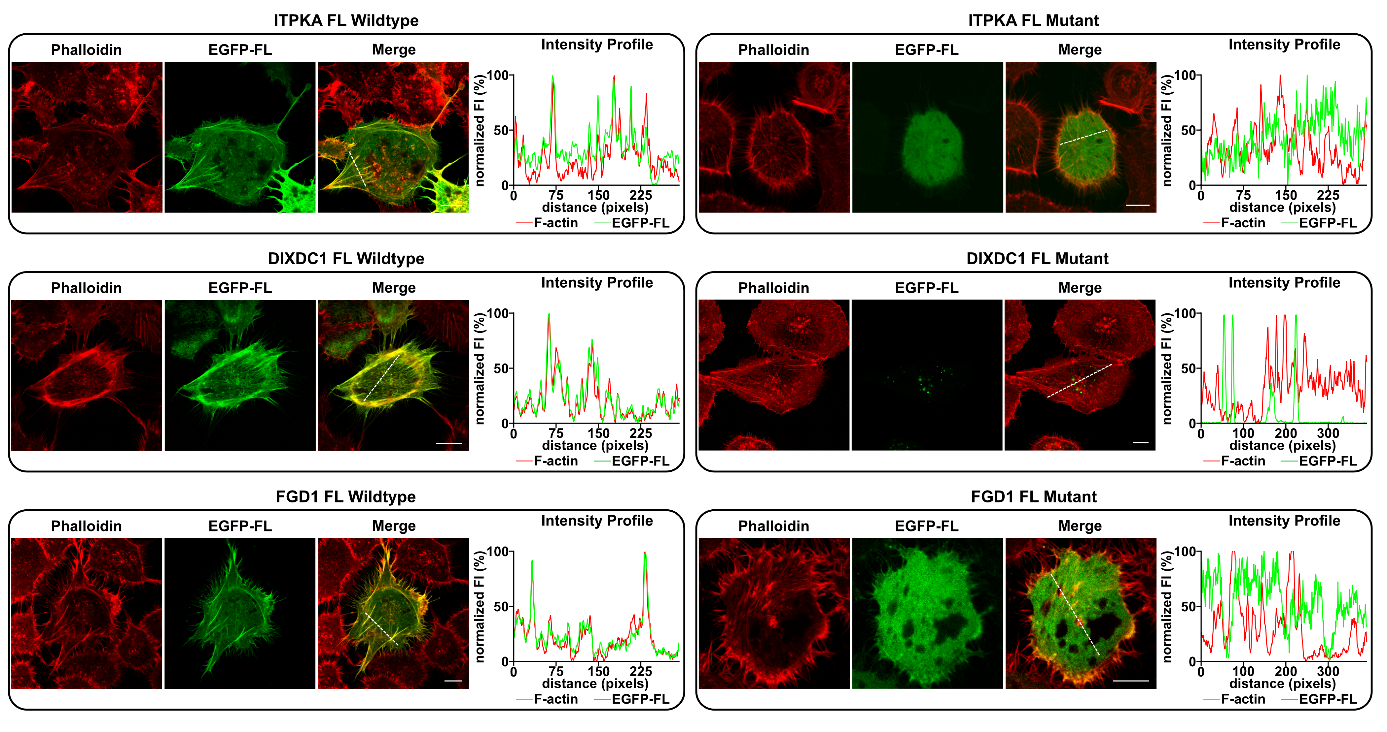
 **Extended Data Figure 6: Colocalization of phalloidin-stained actin and EGFP-tagged full-length proteins in H1299 cells.** Representative confocal images show H1299 cells transfected with wild‐type or mutant full‐length (FL) constructs fused to EGFP (green), together with phalloidin‐568 staining of F‐actin (red). Merged panels highlight varying extents of colocalization (appearing yellow). Intensity profiles (right panels) were recorded along the dashed white lines in each cell, quantifying relative fluorescence for F‐actin and the respective EGFP‐FL protein. Wild‐type constructs generally exhibit robust colocalization with actin filaments, whereas alanine substitutions at key motif positions (P1 or P8) disrupt or reduce F‐actin association. Scale bars, 10 µm.

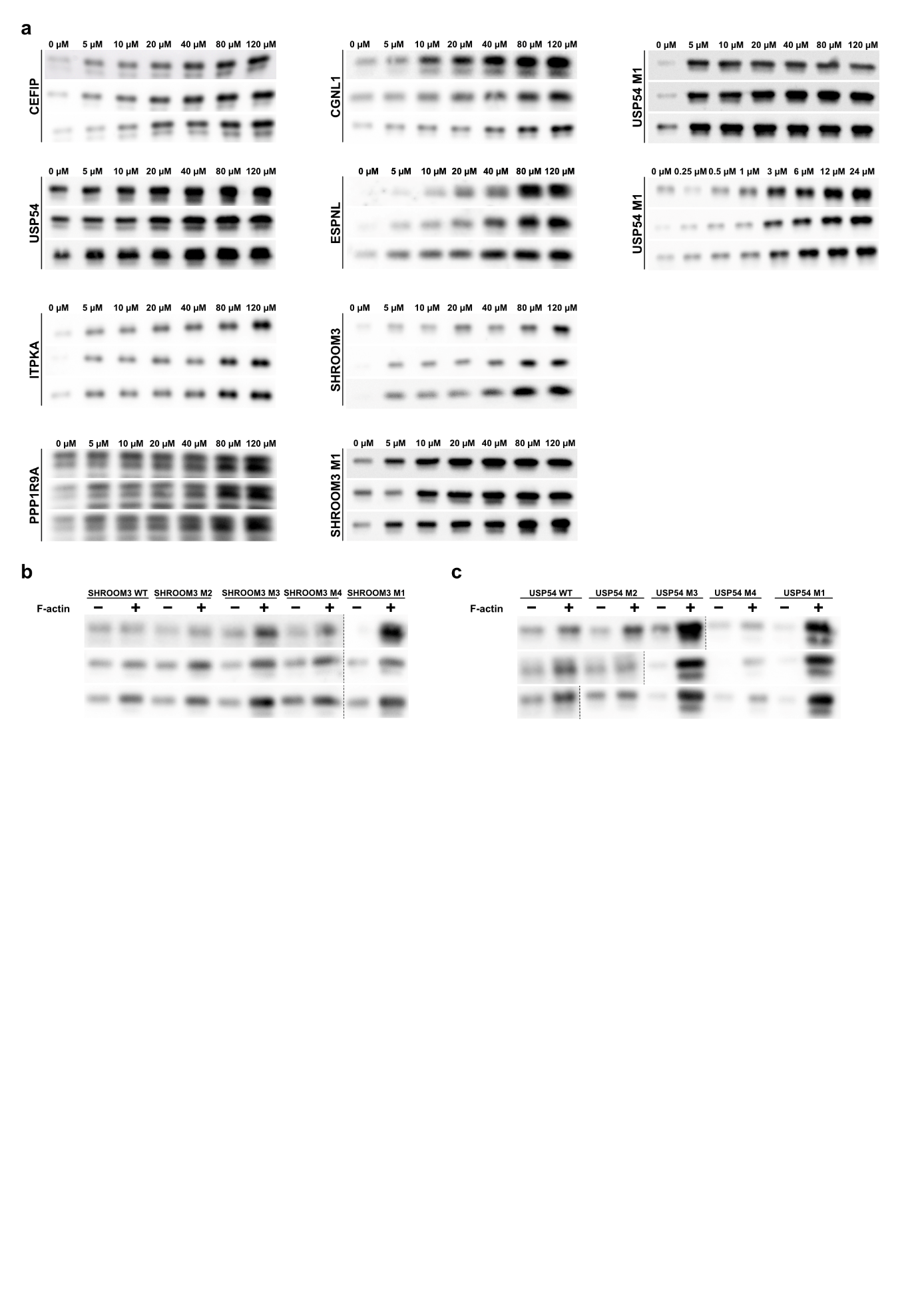
**Extended Data Figure 7: In vitro F-actin pulldown assay of EGFP-tagged peptides and mutant analyses.** **(a)** Representative Western blots of F-actin pulldown assays for each peptide. The pellet fraction is shown, with peptide bands visualized using an anti-EGFP antibody. These assays were performed to determine the binding affinities (K_D_) of wild-type (WT) and mutant peptides. **(b)** Quantified relative band intensities from SHROOM3 pulldown assays, normalized and averaged across replicates, showing differences in binding affinities between WT and mutants. **(c)** Quantified relative band intensities from USP54 pulldown assays, normalized and averaged across replicates, highlighting differences in binding affinities between WT and mutant peptides.

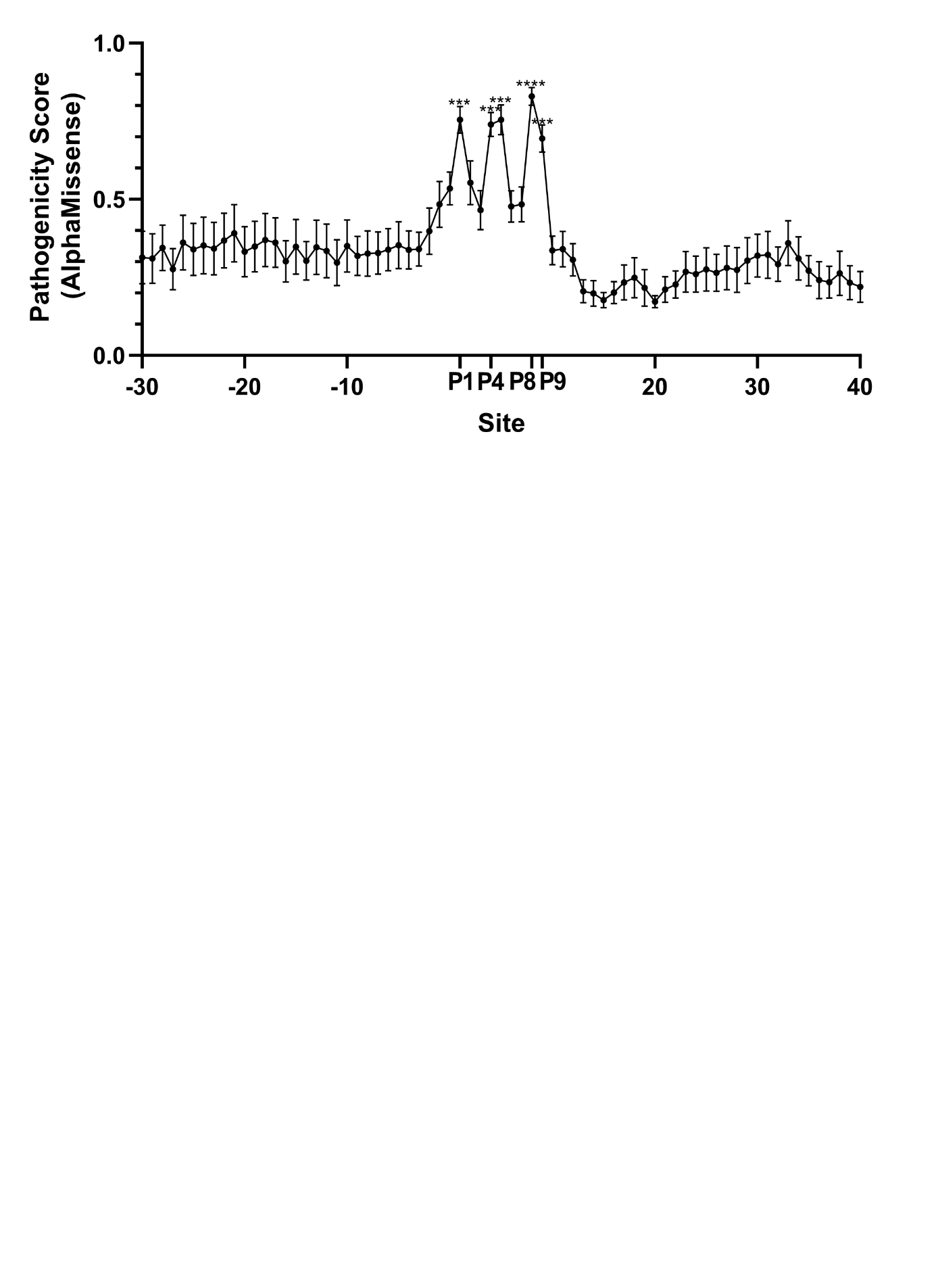
**Extended Data Figure 8: AlphaMissense pathogenicity scores for the conserved SFM region.** Mean pathogenicity scores ± SEM are shown for all peptides tested in this study (both *in cellulo* and *in vitro*). Key motif positions (P1, P4, P5, P8 and P9) exhibit significantly higher AlphaMissense scores than control regions (positions −30 to −20 and 30 to 40), as determined by one‐way ANOVA with multiple comparisons. Asterisks indicate levels of statistical significance (*p < 0.05, **p < 0.01, ***p < 0.001, ****p < 0.0001), consistent with the low dN/dS ratios (see Figure 6c) observed in the same motif region.

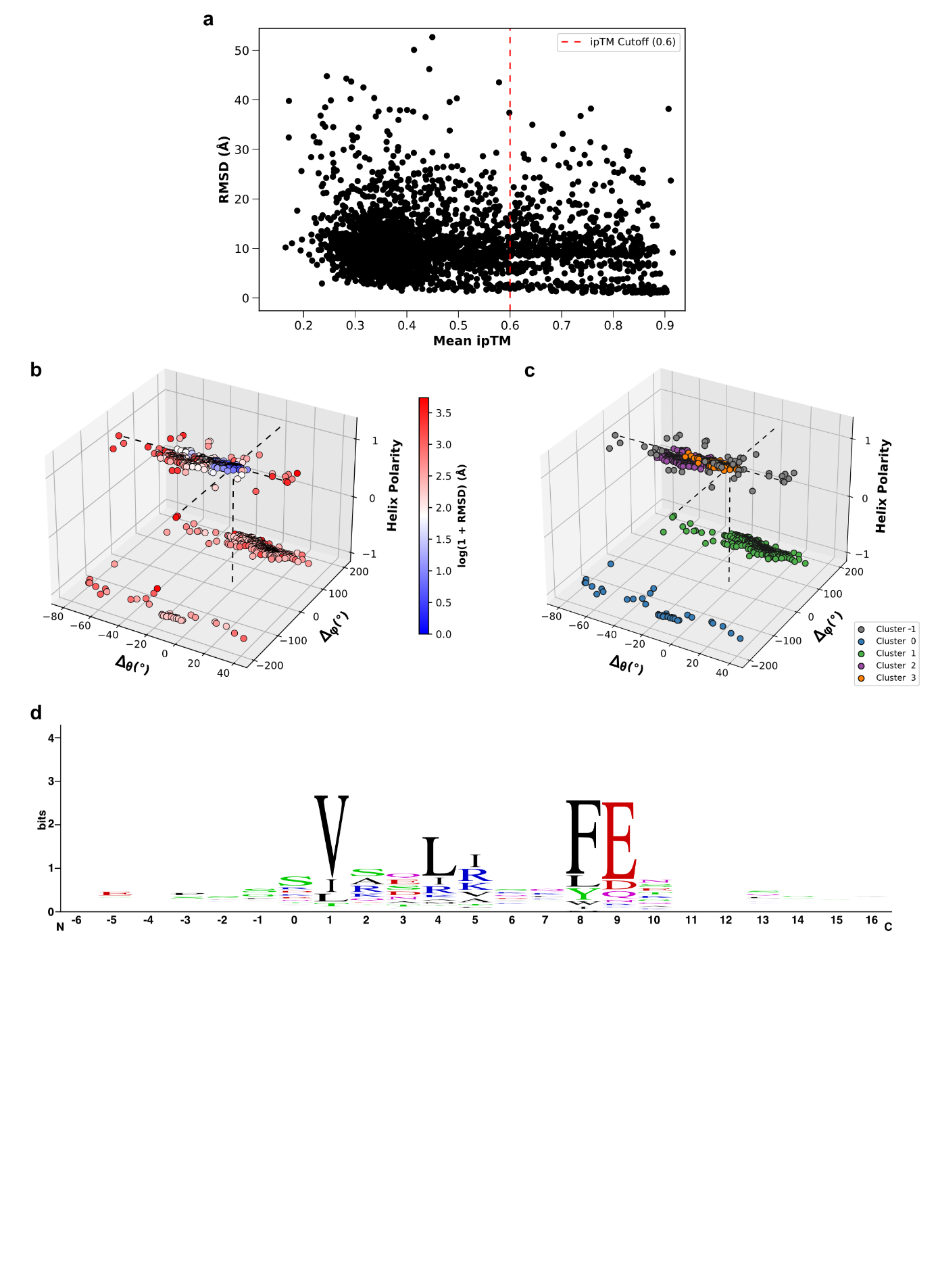
**Extended Data Figure 9: Large‐scale SLiMFold pipeline results from all three iterations (3,414 total AF2 predictions). (a)** Scatterplot of predicted peptide–actin complexes (n=3,414) displaying RMSD relative to the ITPKA reference motif (y‐axis) vs. mean ipTM score (x‐axis). The dashed red line at ipTM=0.6 indicates the threshold below which predicted interfaces are typically deemed unreliable. **(b)** 3D plot of Δφ, Δθ, and helix polarity for each prediction, colored by log(1+RMSD), illustrating the orientation distributions of all modeled peptide–actin complexes. **(c)** The same 3D orientation space as in (b) but color‐coded by HDBSCAN clusters (0–3) and outliers (−1). Cluster 3, in particular, exhibits minimal RMSD and orientations similar to ITPKA. **(d**) Sequence logo of cluster 3, highlighting the core SFM residues (P1, P4, P5, P8 and P9). The logo underscores the high sequence conservation in these positions, consistent with strong structural similarity to the ITPKA–actin reference.

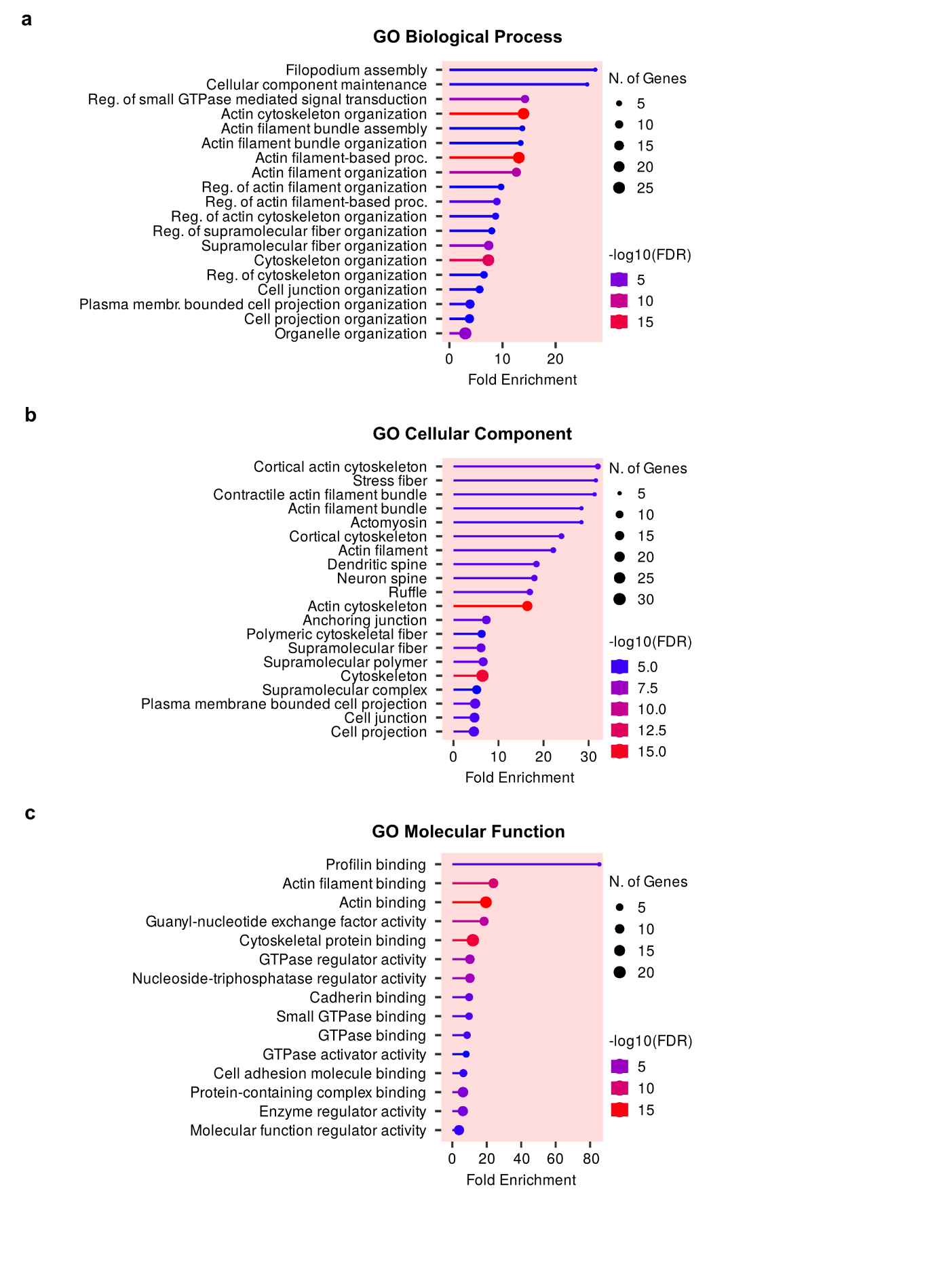
**Extended Data Figure 10**: **Gene Ontology (GO) enrichment analysis of all motif-carrying genes identified by the SLiMFold pipeline.** Shown are the enriched **(a)** biological processes, **(b)** cellular components, and **(c)** molecular functions, with fold enrichment on the x-axis. Dot size represents the number of genes in each GO category, and dot color signifies the false discovery rate (FDR).
